# Single-cell transcriptomics characterizes cell types in the subventricular zone and uncovers molecular defects underlying impaired adult neurogenesis

**DOI:** 10.1101/365619

**Authors:** Vera Zywitza, Aristotelis Misios, Lena Bunatyan, Thomas E. Willnow, Nikolaus Rajewsky

## Abstract

Neural stem cells (NSCs) contribute to plasticity and repair of the adult brain. Niches harboring NSCs are crucial for regulating stem cell self-renewal and differentiation. We used single-cell RNA profiling to generate an unbiased molecular atlas of all cell types in the largest neurogenic niche of the adult mouse brain, the subventricular zone (SVZ). We characterized > 20 neural and non-neural cell types and gained insights into the dynamics of neurogenesis by predicting future cell states based on computational analysis of RNA kinetics. Furthermore, we apply our single-cell approach to mice lacking LRP2, an endocytic receptor required for SVZ maintenance. The number of NSCs and proliferating progenitors was significantly reduced. Moreover, Wnt and BMP4 signaling was perturbed. We provide a valuable resource for adult neurogenesis, insights into SVZ neurogenesis regulation by LRP2, and a proof-of-principle demonstrating the power of single-cell RNA-seq in pinpointing neural cell type-specific functions in loss-of-function models.

**HIGHLIGHTS:**

- unbiased single-cell transcriptomics characterizes adult NSCs and their niche
- cell type-specific signatures and marker genes for 22 SVZ cell types
- Free online tool to assess gene expression across 9,804 single cells
- cell type-specific dysfunctions underlying impaired adult neurogenesis

## INTRODUCTION

Adult neurogenesis is important for the cellular plasticity of the brain. In the adult mammalian brain, the generation of new neurons is restricted to two major sites: the subventricular zone (SVZ) of the lateral ventricles and the subgranular zone (SGZ) in the dentate gyrus of the hippocampus. In the adult SVZ, radial-glia like cells serve as neural stem cells (NSCs) and give rise to transient amplifying progenitors (TAPs), which in turn generate neuroblasts (NBs). NBs migrate along the rostral migratory stream towards the olfactory bulb, where they terminally differentiate into specific subtypes of interneurons and integrate into existing neural circuits (reviewed in Ming and Song, 2011; Figure 1, left).

Adult NSCs reside in a specialized niche, where they are in immediate contact with a variety of different cell types. Cell-intrinsic factors and the microenvironment provided by the niche are both crucial for balancing stem cell self-renewal, proliferation and differentiation (reviewed in Bjornsson et al., 2015). Understanding the potential of single NSCs and the underlying principles of NSC regulation may pave the way toward using these cells as an endogenous source for treating brain injuries or neurodegenerative diseases.

In order to characterize the entire SVZ neurogenic niche at the single-cell level in an unbiased and comprehensive way, we used Drop-seq, a microfluidics- and nanodroplet-based highly parallel transcriptome profiling technology (Macosko et al., 2015). In contrast to recent single-cell RNA sequencing studies, which isolated NSCs and few selected cell types based on the expression of marker genes (Basak et al., 2018; Dulken et al., 2017; Llorens-Bobadilla et al., 2015; Luo et al., 2015; Shin et al., 2015), our strategy circumvents pre-selection. Thereby, we include all SVZ residing cells and minimize the risks of missing potentially undescribed or marker negative cell populations. We identified more than 20 neural and non-neural cell types residing in the SVZ and provide their gene expression signatures. Sequencing data are readily accessible via a freely accessible online tool that allows investigators to evaluate and visualize the expression of their genes of interest in the context of 22 distinct cell types and 9,804 individual cells. We resolved NSC activation states and uncovered differences in RNA dynamics in NSCs and their progeny by estimating future cell states based on changes in mRNA splicing kinetics (La Manno et al., 2017).

We used the unbiased single-cell transcriptome profiling approach to investigate cell-type specific molecular changes in the SVZ of mice mutant for low-density lipoprotein receptor-related protein 2 (LRP2, also known as megalin). LRP2 is a multifunctional endocytic receptor expressed in multiple tissues including the SVZ of the adult brain. LRP2 activity is indispensable for proper development and function of various tissues, including forebrain, eye and heart (reviewed in Christ et al., 2016). In humans, mutations in *LRP2* are the cause of Donnai-Barrow syndrome (Kantarci et al., 2007), a severe autosomal recessive disorder affecting multiple organ systems. These abnormalities are recapitulated in LRP2-deficient mice (Willnow et al., 1996). With relevance to our study, LRP2 is expressed in the ependymal cell layer facing the neural stem cell niche in the SVZ. Loss of LRP2 expression in gene-targeted mice results in impairment of adult neurogenesis specifically in the SVZ, but not the SGZ (where the receptor is not expressed) (Gajera et al., 2010). To investigate which cells types in the SVZ niche are affected and how, we performed comparative single-cell RNA sequencing of SVZ tissues dissected from LRP2-deficient mice and matched littermate controls. We show that the number of NSCs and proliferating cells is reduced in mutant mice and that TAPs express fewer genes associated with cell cycle control. Differential gene expression analysis per cell type revealed perturbations in BMP4 and Wnt signaling in NBs and TAPs, respectively, and a global reduction in ribosomal gene expression. We validated these single-cell RNA sequencing results using immunohistochemistry. Based on our data, we propose a role for LRP2 in integrating the activity of several morphogen pathways in the SVZ to provide the appropriate microenvironment for functional neurogenesis to proceed.

Taken together, this study demonstrates that our unbiased single-cell RNA sequencing strategy enables an untargeted analysis of NSCs within the context of the entire neurogenic niche, reveals consequences of LRP2 ablation on adult SVZ neurogenesis, and provides a resource for studying adult neurogenesis.

## RESULTS

### The entire adult subventricular zone neurogenic niche at single-cell resolution

To study adult neural stem cells, their progeny and surrounding niche cells comprehensively and in an unbiased manner, we microdissected the SVZs from three to five mice, generated a high quality single-cell suspension using an optimized tissue dissociation and debris removal protocol (see experimental procedures for more details), and performed Drop-seq (Macosko et al., 2015), a droplet based single-cell transcriptome profiling method to simultaneously obtain the transcriptome of thousands of cells (Figure 1). Computational analysis of sequencing data was performed with Seurat v1.4 (Satija et al., 2015). To reduce the dimension of data, the most highly variable genes were selected and used for principal component (PC) analysis. All significant PCs were then selected for clustering with the shared nearest neighbor (SNN) algorithm and for data visualization with t-distributed stochastic neighbor embedding (tSNE) as implemented in Seurat.

**Figure 1.**
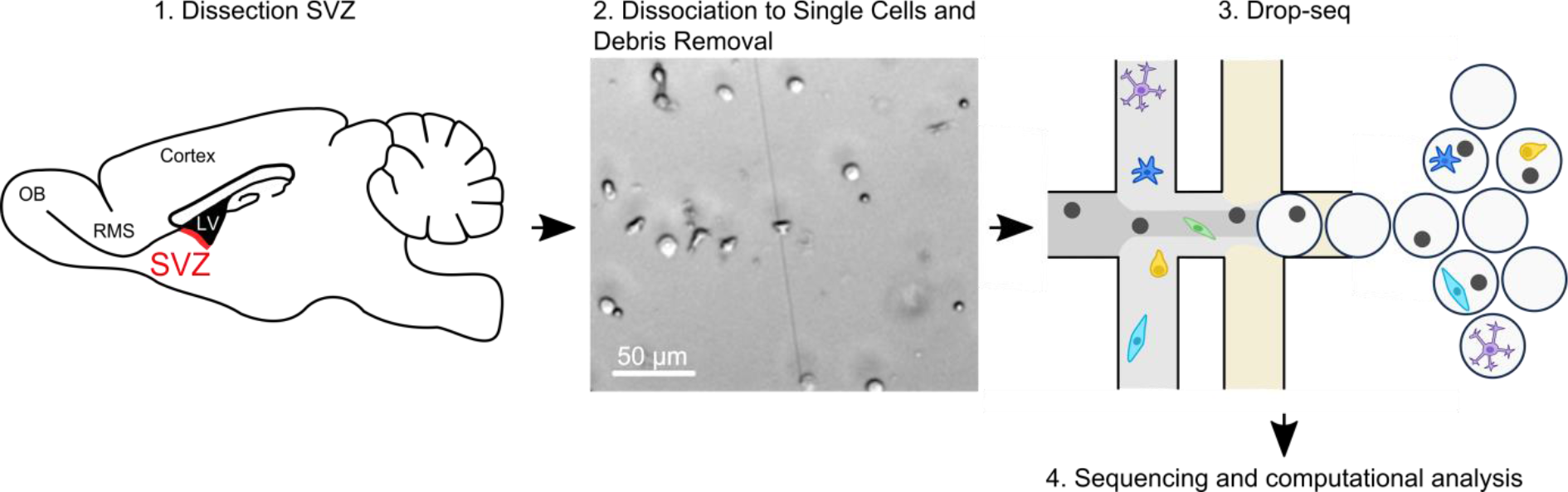
Single-cell RNA sequencing of the adult SVZ. After microdissection of the SVZ (1, marked in red), the tissue was dissociated into single cells. Debris, dead cells and cell clumps were removed. The single-cell suspension (2) is applied to a microfluidics device, where single cells are combined with barcoded beads in droplets (3). After library preparation and sequencing, single-cell transcriptomes were deconvolved and analyzed (4). left: sagittal section view of the adult mouse brain.

**Abbreviations:** SVZ: subventricular zone; OB: olfactory bulb; RMS: rostral migratory stream; LV: lateral ventricle.

We collected 9,804 cells with 734 genes and 1,137 UMIs quantified per cell (medians) in five independent Drop-seq experiments (Figure S1A). Independent analysis of replicates revealed similar results (data not shown). To gain higher resolution, we analyzed cells from all replicates jointly. We did not observe batch effects as cell clusters contained cells from all replicates in similar proportions (Figures S1A and S1B). Initial analysis identified 17 distinct cell clusters (Figure 2A), which could be assigned to known cell types based on the detection of known marker genes (Figures 2B, S2 and Table S1). Cell type enriched genes were identified by comparing each cluster to all others (Table S1). Exemplarily, we show the expression of the top five genes from each cell cluster across all cells (Figure 2C).

### Characterization of cell types residing in the adult SVZ

NSCs formed a cluster in the center of the tSNE plot (Figure 2A). We identified them based on (1) their similarity to astrocytes (Doetsch et al., 1999), (2) the expression of *Thbs4*, which was previously reported to be highly enriched in NSCs (Beckervordersandforth et al., 2010; Llorens-Bobadilla et al., 2015), and (3) upregulation of genes associated with NSC activation such as *Ascl1* at the upper part of the cluster (Figure 2D, blue arrow). The commonly used stem cell markers *Nes* and *Prom1* (CD133 in human) (reviewed in Chaker et al., 2016) were rarely expressed in NSCs, but were mostly detected in endothelial cells (Figure S3, see below). NSCs shared the expression profile with astrocytes. For example, *Slc1a3* (*Glast*) was highly enriched in both cell types. By contrast, NSCs and most astrocytes were negative for *S100b* (here mainly detected in MOLs and ependymal cells) (Figure 2D), which is a marker for astrocytes residing deeper in the tissue at the interface to the striatum (Codega et al., 2014). These findings strongly suggest that cluster 17 and 16 of our dataset consists of NSCs and niche astrocytes, respectively. Despite overlapping expression profiles, the breadth of our dataset enabled us to resolve NSCs from niche astrocytes and to identify significantly up- and downregulated genes (Table S2). In Figure 2D, we show exemplarily that *Aqp4*, a marker of mature astrocytes, is barely detected in NSCs (green circle). Notably, previous scRNA-seq studies could not separate NSCs and niche astrocytes (Basak et al., 2018; Dulken et al., 2017).

**Figure 2.**
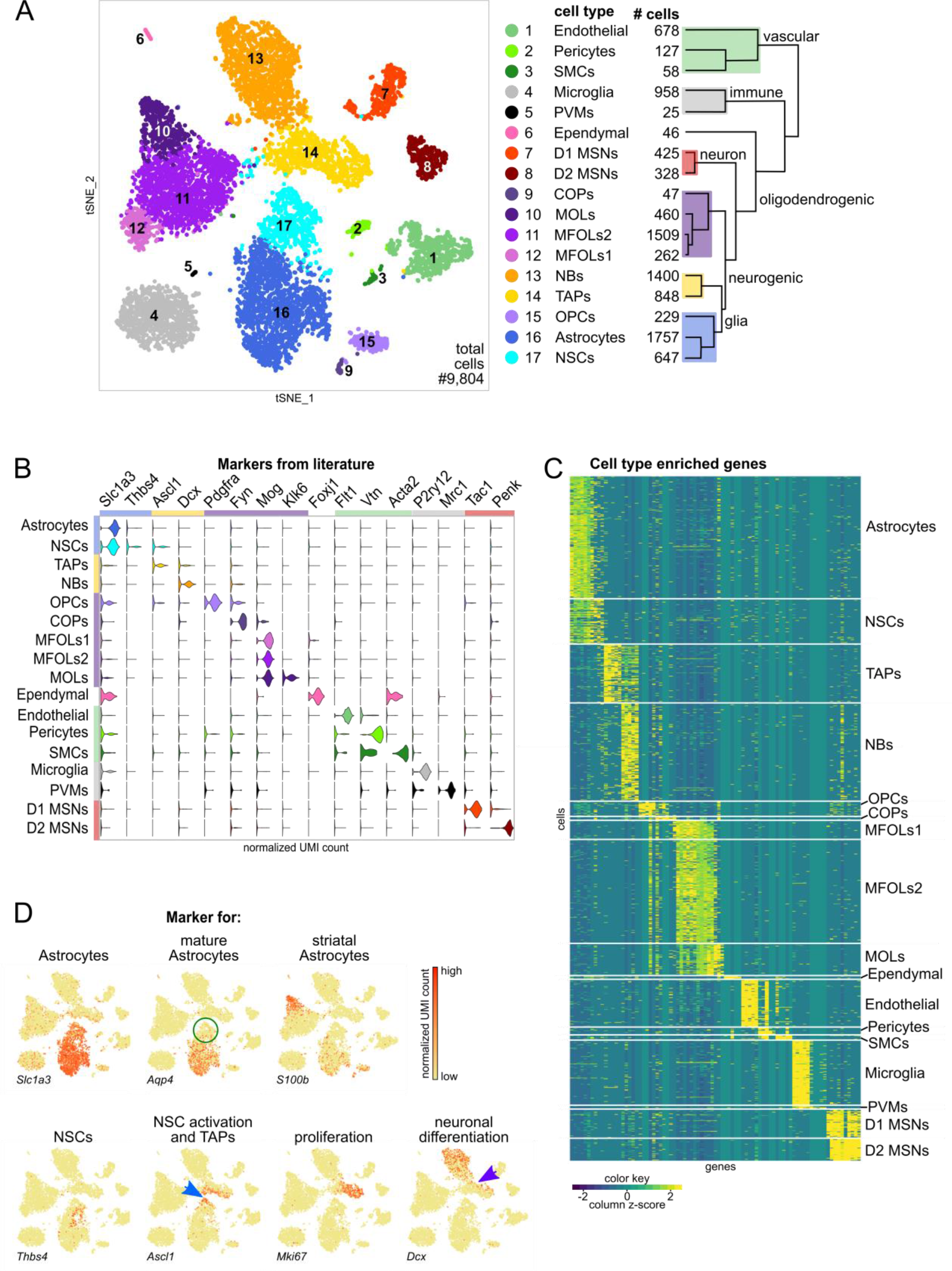
Characterization of cell types residing in the adult SVZ. (A) tSNE plot of 9,804 cells colored by cluster annotation. The dendogram displays the relationships of cell clusters. Underlying boxes highlight main cell classes. (B) Identification of cell types based on known marker genes. See also Figures S2 and S3. (C) Heatmap depicting the expression of the top five enriched genes per cell type across all identified cells. Each row represents a single cell and each column represents a gene. The expression is normalized by gene. For all significantly upregulated genes per cell type see Table S1. (D) tSNE plots of cells colored by expression of selected marker genes, which were used for the identification of astrocytes, NSCs, TAPs and NBs. The color key indicates expression levels (red: high, yellow: low). Green circle highlights absence of *Aqp4* in NSCs. Blue arrow indicates expression of *Ascl1* in NSCs, which are close to TAPs. Purple arrow points to TAPs expressing *Dcx*.

**Abbreviations**: SMCs: smooth muscle cells; PVMs: perivascular macrophages; MSNs: medium spiny neurons, which are contaminants from the striatum; COPs: differentiation-committed oligodendrocyte precursors; MOLs: mature oligodendrocytes; MFOLs: myelin forming oligodendrocytes; NBs: neuroblasts; TAPs: transient amplifying progenitors; OPCs: oligodendrocyte progenitor cells; NSCs: neural stem cells.

Adjacent to NSCs, TAPs composed a cluster enriched for proliferation markers such as *Mki67* and *Pcna*, and genes associated with neuronal commitment e.g., *Dlx1* and *Dlx2* (Figures 2D and S2). We detected genes known to be involved in neuronal differentiation (e.g., *Dcx*, *Tubb3*) in TAPs, which are close to the NB cluster (Figure 2D, purple arrow). The expression of neuronal marker genes increased towards the tip of cluster 13 (NBs) (Figures 2D and S2).

Based on the expression of marker genes such as *Flt1*, we identified cells of cluster 1 as endothelial cells (Figures 2B and S3). As mentioned before, we observed that these cells express markers associated with other cell types in the literature. For example, PROM1 (CD133 in human), a microvilli and primary cilia associated protein (Dubreuil et al., 2007; Weigmann et al., 1997), is widely used to identify and isolate ependymal cells, and, in combination with GFAP or GLAST, NSCs (Beckervordersandforth et al., 2010; Fischer et al., 2011; Llorens-Bobadilla et al., 2015). At the RNA level, we detected *Prom1* to be co-expressed with known markers of endothelial cells (e.g., *Flt1*, *Slc2a1/Glut1*) (Figures S3A and S3B). In addition to *Prom1*, endothelial cells were positive for *Vim* and *Nes*, markers for ependymal and neural progenitor cells, respectively (Figures S3A and S3B). Vice versa, we detected the endothelial marker *Slc2a1* to be co-expressed with the ependymal markers *Foxj1* and *Ak7* in cluster 6, and therefore conclude that endothelial and ependymal cells share the expression of some established marker genes (Figures S2, S3A, and S3B). Evaluation of (1) *in situ* images of the Allen Brain Atlas (Lein et al., 2007), and (2) bulk RNA-seq data from purified cell populations (Zhang et al., 2014) supports our findings (Figures S3C and S3D).

Furthermore, we identified, two types of mural cells (pericytes and smooth muscle cells, short SMCs), two distinct clusters consisting of immune cells (microglia and perivascular macrophages, short PVMs), five clusters comprising different stages of the oligodendrocyte lineage (oligodendrocyte progenitor cells, short OPCs; differentiation-committed oligodendrocyte precursors, short COPs; myelin forming oligodendrocytes, short MFOLs1 and MFOLs2; mature oligodendrocytes, short MOLs; Marques et al., 2016), and two clusters of mature neurons (markers used for cell type identification in Figures 2B and S2). Notably, OPCs clustered close to NSCs and astrocytes, indicating higher similarity of OPCs to astroglia than oligodendroglia (Figure 2A dendogram).

Based on the expression of marker genes like *Tac1* and *Pdyn* or *Penk* and *Adora2a* we conclude that the observed mature neurons are D1 and D2 MSNs, respectively (Figures 2B and S2) (Gokce et al., 2016), which likely originate from the underlying striatum. Recently, it has been shown that acetyltransferase (ChAT)-positive neurons reside in the SVZ (Paez-Gonzalez et al., 2014). However, the only known marker for this SVZ residing neuron type (*Chat)* was not detected in our dataset. We observed that a few cells of cluster 7 separated in the tSNE space and were negative for D1 and D2 MSNs markers (Figure S2, red circle) indicating that they are a distinct neuron type and may represent SVZ residing ChAT neurons.

Taken together, our unbiased single-cell RNA profiling strategy enabled the explicit identification of cell types based on a multitude of genes, gives an estimate on their relative proportions in the SVZ, and characterizes the majority of neural and non-neural cell populations known to constitute the adult SVZ stem cell niche (for review see Bjornsson et al., 2015; Bonaguidi et al., 2016) at unprecedented resolution. We provide a user-friendly online tool (https://shiny.mdcberlin.de/SVZapp/) for easy access to our data allowing visualization of gene expression in 9,804 individual cells and in the context of distinct cell types.

### RNA dynamics reveal heterogeneity within the neurogenic lineage

To gain insight into the dynamics of stem cell activation and differentiation in the neurogenic lineage, we used velocyto (La Manno et al., 2017), a computational method that predicts the future state of individual cells from single-cell transcriptome data. The underlying assumption behind velocyto is that changes in the transcriptional rate of a gene can be used to predict its future expression level. For each gene, these changes can be estimated from the ratio of unsliced to spliced reads. Calculating the RNA velocity of the most variable and expressed genes in a single cell results in the estimated future expression state, which the cell is about to accomplish in a timescale of a few hours (La Manno et al., 2017).

We used velocity to investigate the relationship between astrocytes, NSCs, TAPs, and NBs (Figure 3A). We visualized the results in tSNE space by plotting an arrow for each cell, which spans the actual and the estimated future state. The arrow length reflects the magnitude of the calculated velocity. Hence, cells undergoing vigorous transcriptomic changes have long arrows, whereas cells with little RNA dynamics have short to no arrows. In the astrocyte cell cluster, we observed little and uncoordinated RNA velocity indicating that these cells were in a transcriptionally stable state undergoing few changes. NSCs separated from the astrocyte cluster demonstrating distinct transcriptome signatures. We observed differences in RNA dynamics within the NSC and neural progenitor clusters. RNA velocity was low in NSCs close to astrocytes, increased towards TAPs, and decreased in NBs. To gain higher resolution of cell types constituting the neurogenic lineage, we performed subclustering of NSCs, TAPs, and NBs (cluster 17, 14, 13, respectively, in the all cell tSNE) and obtained eight clusters, six of which represent different progenitor states along the neurogenic lineage (Figure 3C). Subcluster 1 was composed of contaminating T cells, whereas subcluster 2 contained oligodendrocyte-like cells (description in Figures S4C-E). Subclusters were composed of cells from all replicates with only slight differences in ratios in three out of eight subclusters (Figures S4A and S4B). Subcluster enriched genes are provided in Table S3.

To classify the six subclusters of the neurogenic lineage, we evaluated the expression of marker genes and scored the cells based on previously published gene sets (Llorens-Bobadilla et al., 2015) (Figure 3D). We identified quiescent NSCs (qNSCS) based on highest expression levels of genes associated with lipid biosynthesis, glycolysis, and glial markers. In activated NSCs (aNSCs) transcript levels of cell cycle-associated and ribosomal genes started to increase and rose further in TAPs. Genes associated with mitosis were almost exclusively detected in subcluster 5. Therefore, we designated these cells mitotic TAPs (mTAPs). In NBs, the expression of genes associated with cell cycle and mitosis decreased while the expression of genes indicative of neuronal differentiation increased (Figure 3D). NBs formed two clusters and neuronal genes were more lowly expressed in early compared to late NBs. To gain insight into the progression of cell cycle, we visualized cells within S and G2M phase by summarizing expression values of respective genes from an independent gene list (Kowalczyk et al., 2015) (Figures S4G and S4H). The tSNE feature plots illustrated the entry of aNSCs into S phase, followed by TAPs, which showed high expression of S phase genes. S phase genes decreased towards the tip of the mTAPs cluster: Here G2M genes displayed highest expression.

The velocity analysis revealed that NSCs have little RNA dynamics during quiescence but change their transcriptome vigorously within few hours upon activation (Figure 3A). The majority of arrows pointed towards the NB cluster and we observed little flow of aNSCs in the opposite direction. While this is in line with the theory that aNSCs do not divide asymmetrically, but mostly undergo consuming (=unidirectional) cell divisions to generate TAPs (Obernier et al., 2018), the knn-smoothing of the velocity analysis might hide outliers going the opposite direction. Arrows within TAPs either pointed towards NBs or to the bottom of the mTAP cluster. mTAPs displayed highest velocity most likely resulting from their fast cycling nature. The vast majority of arrows in the NB cluster pointed away from NSCs and TAPs. Decreasing RNA dynamics towards late NBs made us speculate that cells in this differentiation state have a stable transcriptome in the SVZ, but change their gene expression in the olfactory bulb where they terminally differentiate.

We identified the root of the differentiation process in the NSC cluster (Figure 3B). In conclusion, our RNA dynamics analysis independently confirmed the marker-based identification of NSCs and the direction of their differentiation fate via TAPs to NBs. Together with the subclustering analysis, we gained insights into the transcriptional dynamics of different NSC activation states and progeny subtypes. The continuity of cells in the tSNE space together with the progression of gene expression along cell types (Figure 3D) (1) demonstrates that our dataset contains the entire neurogenic lineage, and (2) illustrates the continuous nature of neurogenesis which makes a rigid classification of distinct cell types along the differentiation trajectory difficult.

### Identification of NSC activation stage enriched genes

The comprehensiveness of our datasets enables evaluation of expression levels of cell type enriched genes in the context of the entire niche. At the moment, the discrimination of astrocytes, qNSCs, aNSCs, and TAPs based on single marker genes is challenging. Manual inspection of cluster enriched genes (Tables S2 and S3) revealed several candidates, which might be involved in stem cell regulation and could be used to better discriminate NSCs from astrocytes and TAPs (Figure S4F). For example the long non coding RNA *Meg3* is known to function as a negative regulator of growth (Zhou et al., 2012). In our dataset, it was highly expressed in mature neurons, but also detected in NSCs, NBs, OPCs, and ependymal cells. Surprisingly, it was absent from astrocytes and inversely correlated with genes associated with cell cycle as well as mitosis (Figures S4E blue background, and S4G-I). It would be interesting to investigate, why *Meg3* is expressed in NSCs, but not in astrocytes. We hypothesize that it is involved in maintaining quiescence and preventing cells from entering the cell cycle.

**Figure 3.**
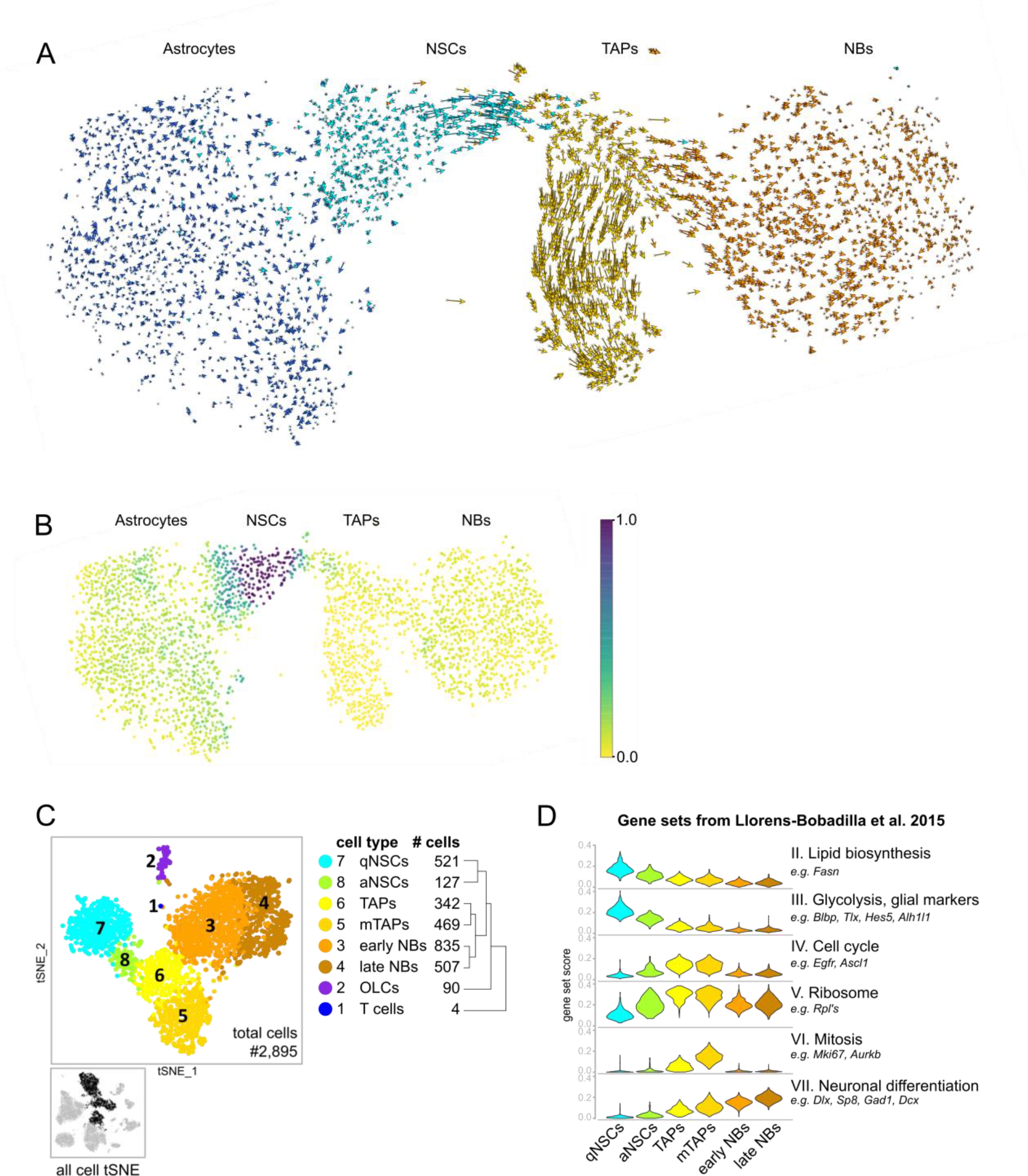
RNA dynamics reveal heterogeneity within the neurogenic lineage. (A) RNA velocity plotted in tSNE space for astrocytes, NSCs, TAPs, and NBs. For each cell, arrows show the location of the estimated future cell state. RNA dynamics differ between cell clusters and within NSCs, TAPs, and NBs. (B) The root of the differentiation process was identified in the NSC cluster by modeling the transition probabilities derived from the RNA velocity. The color scale represents the density of starting points of a reverse Markov process and ranges from low (yellow) to high (blue). (C) Subclustering analysis of NSCs, TAPs, and NBs (marked in black in the original tSNE in the lower left) revealed eight subclusters. The dendogram shows the relationships of cell types. Subcluster 1 and 2 are described in Figures S4C-E. Subclusters 3-8 belong to the neurogenic lineage and can be recognized by their RNA dynamics in the velocity tSNE depicted in (A). (D) Cell type characterization based on gene sets published in (Llorens-Bobadilla et al., 2015).

**Abbreviations:** NSCs: neural stem cells; TAPs: transient amplifying progenitors; NBs: neuroblasts. qNSCs: quiescent NSCs, aNSC: activated NSCs, mTAPs: mitotic TAPs; OLCs: oligodendrocyte-like cells.

### single-cell RNA profiling of LRP2-deficient SVZ reveals a reduction of NSCs and proliferating cells

We wondered whether an unbiased transcriptomics approach would be able to uncover cell type-specific molecular defects underlying disturbances in adult SVZ neurogenesis. To do so, we used mice genetically deficient for expression of LRP2. LRP2, a member of the low-density lipoprotein receptor gene family, acts as an endocytic retrieval receptor for several signaling molecules and morphogens, including bone morphogenetic protein (BMP) 4 and sonic hedgehog (SHH) (Christ et al., 2012, 2015; Gajera et al., 2010). It plays central roles in controlling embryonic and adult neurogenesis (reviewed in Christ et al., 2016). With relevance to this study, LRP2 is highly expressed on the apical surface of ependymal cells in the adult SVZ, but not detected in the hippocampal neurogenic niche (SGZ) (Gajera et al., 2010). Adult LRP2-deficient mice display mild forebrain formation defects such as enlarged lumens of the lateral ventricles, but the ventricular system and ependymal architecture appeared normal in histological studies (Gajera et al., 2010). Interestingly, the number of NSCs and their progeny is significantly reduced, and neurogenesis is impaired specifically in the SVZ of adult LRP2-deficient mice (Gajera et al., 2010).

To elucidate which cell types are affected by ablation of LRP2 and to gain insights into the underlying molecular causes with cell type resolution, we performed single-cell RNA profiling on SVZ dissected from LRP2-deficient mice (KO) and littermate controls (Ctr). We collected 8,056 cells in two replicates (Ctr and KO, four Drop-seq runs, Figure S5A). Cell cluster proportions and relationships between cell types were similar for individual replicates (data not shown). Joint analysis of all samples resulted in the identification of 18 cell types (Figures 4A and S5B). Overall, we did not observe discernable differences in cluster proportions based on batch or genotype (Figures S5A and S5C), indicating no overt changes in SVZ architecture of the mutant as compared to the control SVZ. Using a generalized linear mixed model to test biological variation independent from technical noise by modeling the genotype as fixed and the batch as random effect, we revealed a significant reduction in NSC proportions (p-value < 0.001, Figure S5C right panel) in agreement with earlier observations obtained by targeted immunohistochemical approaches (Gajera et al., 2010).

We compared the LRP2 KO/Ctr data (Dataset B) with our previous results (Dataset A, see Figure 2) and observed only mild differences. Pericytes and SMCs were combined in one cluster (called “mural cells”) whereas mature neurons formed three clusters comprising potential ChAT neurons, D1, and D2 MSNs. As in Dataset A, potential ChAT neurons expressed markers of mature neurons, but markers of striatal neurons were not detected. Visual inspection of genes discriminating these neurons from D1 and D2 MSNs revealed five candidates, which could serve as new markers (*Kit*, *Nxph1*, *Pthlh*, *Crtac1*, *Sparcl1*) for this particular cell type (Figure S5D). Furthermore, we detected OLCs in an individual cluster and one cluster comprising choroid plexus cells (Figure 4A and S5B). As in Dataset A, subclustering of NSCs, TAPs, and NBs resolved the different stages of neurogenesis (Figure S6). The similarity of Dataset A and B demonstrates the robustness of our method making it suitable to obtain and compare data from different mouse strains and genotypes.

Cells in the adult SVZ proliferate less in LRP2-deficient mice (Gajera et al., 2010). To analyze which cell types are affected, we scored cells according to cell cycle using a list of cell cycle-associated genes (Kowalczyk et al., 2015) and plotted the cumulative fraction distributions per genotype and cell cluster. We quantified significantly less expressed cell cycle genes in TAPs derived from mutant mice as compared to control cells (p-value <0.01, Figures 4B and S7A). Notably, the overall number of TAPs in our sequencing data did not change indicating that LRP2-deficient mice have the same amount of progenitors, but specifically fast cycling TAPs divide less. To quantify fast proliferating cells in the SVZ, we injected LRP2-deficient mice and littermate controls by a single intraperitoneal injection of BrdU and sacrificed the animals 24 hours later. Counting of BrdU positive cells per ventral SVZ section confirmed a significant reduction of fast-proliferating cells in LRP2-deficient mice (Figures 4C and 4D).

**Figure 4.**
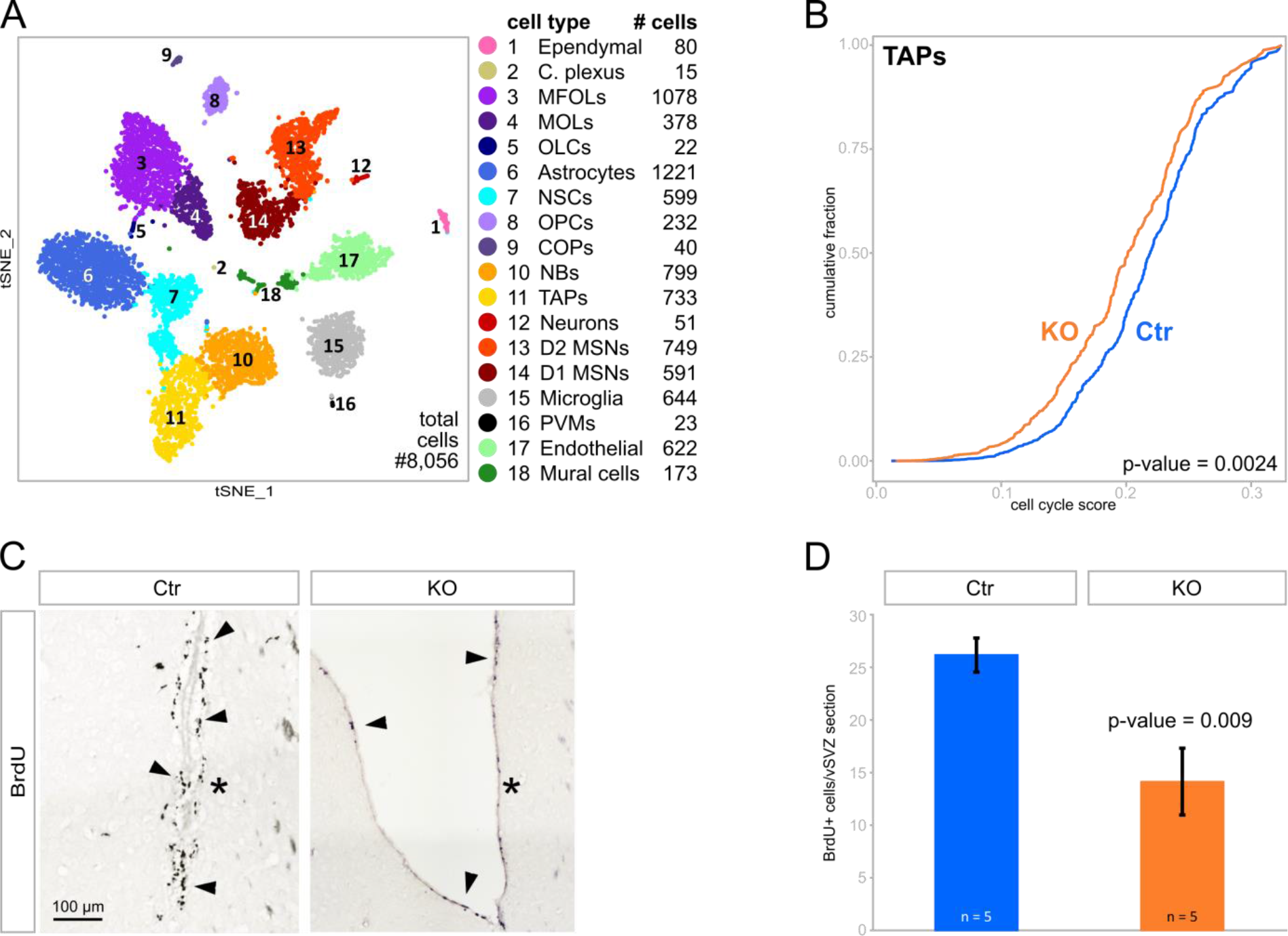
Cell proliferation is reduced in the SVZ of LRP2-deficient mice. (A) tSNE plot of 8,056 cells from the SVZ of LRP2-deficient mice and littermate controls, colored by cluster annotation. For cell type identification see Figure S5B. (B) Cumulative fraction of LRP2 KO and Ctr TAPs scored for cell cycle. Statistical significance of data was determined using Kolmogorov-Smirnov test. (C) Immunohistological detection of BrdU+ cells in the ventral (v) SVZ of KO and Ctr mice. BrdU+ cells are highlighted by arrowheads. Asterisks label the lateral ventricular wall. (D) Quantification of the number of BrdU+ cells in the vSVZ at 24 hours after BrdU injection (as exemplified in panel C). Mean values ± standard error of the mean were determined by counting the number of BrdU+ cells on a total of 12 sections for each animal (five animals per genotype). Statistical significance of data was determined using unpaired student’s t test.

**Abbreviations**: C. plexus: choroid plexus; OLCs: oligodendrocyte-like cells, other cell type abbreviations as in Figure 2; vSVZ: ventral SVZ; KO: LRP2-deficient mice; Ctr: wildtype and heterozygous littermate controls.

### Cell type-specific differential gene expression analysis revealed perturbations in BMP4 and Wnt signaling, and a reduction in ribosomal gene expression in LRP2-deficient mice

To identify cell type-specific molecular changes underlying the impaired adult SVZ neurogenesis in LRP2-deficient mice, we performed differential gene expression analysis per cell cluster. We analyzed the two replicates independently (an007_WT vs. an007_KO, and an010_WT vs. an010_KO) and considered only genes with a p-value smaller than 0.01 and a consistent fold change in both replicates as differentially expressed (Table S4). The results for NSCs, TAPs, and NBs, are displayed in Figures 5A-C. Interestingly, we found *Id3*, a downstream target of BMP4 signaling, upregulated in LRP2-mutant mice, in agreement with earlier findings using semi-quantitative immunohistology (Gajera et al., 2010). *Id3* mRNA was specifically elevated in NBs derived from the LRP2-deficient SVZ (Figure 5C, green box). Increased immunostaining for ID3 protein in mutant mouse brain sections confirmed this result (Figures 5E and 5F). Furthermore, we found Catenin Beta 1 (*Ctnnb1*) to be decreased in TAPs of LRP2-mutant mice (Figure 5B, blue box). The Wnt/beta-catenin pathway is known to simulate NSC proliferation and self-renewal (Qu et al., 2010), but has not been linked to LRP2 activity so far. However, as the numbers of dividing cells and NSCs are decreased in LRP2-deficient mice, it is likely that LRP2 not only interacts with BMP4 and SHH (Christ et al., 2012, 2015; Gajera et al., 2010), but also other morphogens including Wnt that is known to regulate adult neurogenesis. To substantiate a role for LRP2 in Wnt signaling in the SVZ, we crossed the LRP2 mutant strain with a *Tcf/Lef-LacZ* reporter line commonly used to monitor Wnt/beta-catenin signaling activity *in vivo* (Mohamed et al., 2004). Indeed, immunohistochemistry of SVZ sections revealed significantly reduced *Tcf/Lef-LacZ* activity in LRP2-deficient mice as compared to littermate controls (Figure 5D).

Among the downregulated genes in TAPs and NBs obtained from LRP2-mutant SVZ, we observed many genes encoding ribosomal subunits (Figures 5B and 5C, triangles). To investigate if this is a cell type specific or a global trend, we compared the expression of ribosomal genes to the expression of all remaining genes per cell type. In 10 out of the 18 cell types, ribosomal genes were significantly reduced indicating a broad effect of LRP2-deficiency on ribosomal biogenesis in multiple cell types in the niche (Figure S7B). However, TAPs had by far the lowest p-value (Figures 5G and S7C), which is consistent with the observed reduction in cell proliferation. To confirm this result, we exemplarily stained for the ribosomal protein S6-RP and measured the relative signal intensity per SVZ section. We found significant reduction in the amount of S6-RP in the SVZ of LRP2-deficient mice as compared to littermate controls (Figures 5H and 5I).

**Figure 5.**
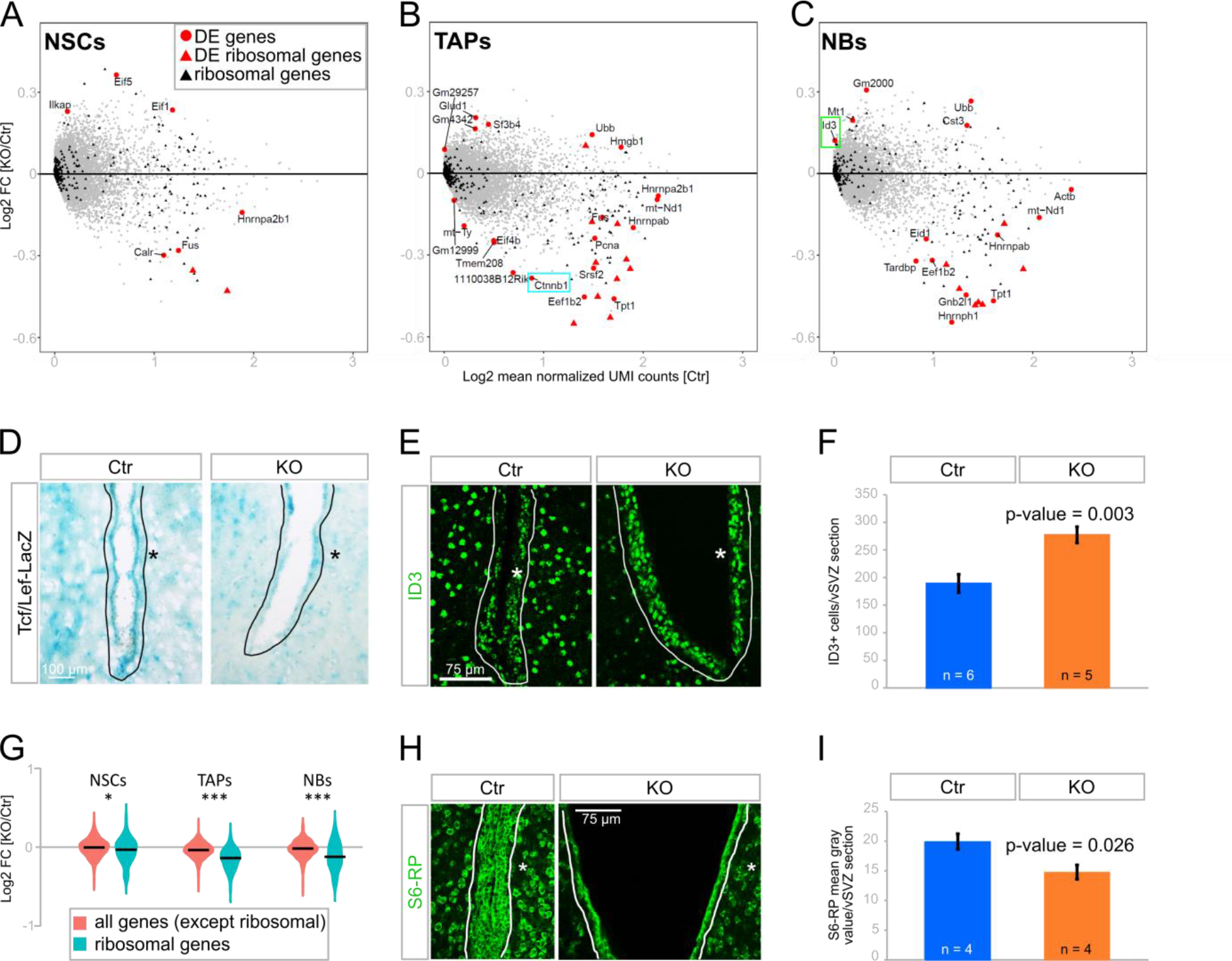
Differential gene expression (DE) analysis per cell type revealed perturbations in BMP4 and Wnt signaling and a reduction of ribosomal genes in LRP2-deficient mice. (A-C) Comparison of gene expression KO vs. Ctr in NSCs, TAPs and NBs, respectively. Ribosomal genes are depicted as triangles. Significantly differential genes are marked in red, non-ribosomal genes are named. Blue box highlights decrease of *Ctnnb1* in TAPs (B), green box highlights increase of *Id3* in NBs (C). Significantly changed genes for all cell types identified in Figure 4A are listed in Table S4. DE analysis was performed using EdgeR. (D) Staining for LacZ activity (blue) on sections of the vSVZ of Tcf/Lef-LacZ mice either Ctr or KO for *Lrp2*. Exemplary images from a total of four mice per genotype are shown (three sections per animal evaluated). (E) Immunohistological detection of ID3 (green) in the vSVZ of Ctr or KO mice. (F) Quantification of the number of ID3+ cells in the vSVZ of Ctr and KO mice (as exemplified in panel E). Three sections each from five KO and six Ctr mice were analyzed. (G) Distribution of KO vs. Ctr expression of ribosomal (red) and all remaining genes (teal) for NSCs, TAPs, and NBs. Expression cut off ≥ 0.5 log2 (mean normalized UMI count) Ctr cells. *p-value < 0.05, *** p-value < 0.001, unpaired student’s t test. For comparison of all cell types see Figure S7B. (H) Immunohistological detection of ribosomal protein S6 (S6-RP, green) in the vSVZ of Ctr or KO mice. (I) Quantification of S6-RP expression levels in the vSVZ of Ctr and KO mice (as exemplified in panel H). Values are given as mean fluorescence intensities. Three sections each from four mice per genotype were quantified. (D, E, H) Solid lines mark the SVZ. Asterisks label the lateral ventricular wall. (F, I) Depicted are mean values +/- standard error of the mean. Statistical significance of data was determined using unpaired student’s t test.

**Abbreviations:** FC: Fold Change; DE: differential expressed; NSCs: neural stem cells; TAP: transient amplifying progenitors, NBs: neuroblasts; KO: LRP2-deficient mice; Ctr: wildtype and heterozygous littermate controls; vSVZ: ventral subventricular zone; S6-RP: ribosomal protein S6.

Taken together, differential gene expression analysis confirmed previously known defects in BMP4 signaling and identified a new signaling pathway (Wnt) to be perturbed in LRP2-deficient mice. In both cases, our analysis was able to newly assign these signaling defects to distinct cell types in the SVZ niche.

## DISCUSSION

The subventricular zone (SVZ) of the lateral ventricles is the largest germinal zone in the adult rodent brain. However, a comprehensive catalogue, that molecularly profiles all cell types residing in this niche, does not exist up to now. In 1997, Doetsch and colleagues published a study addressing the cellular composition of the SVZ. Based on ultrastructure and few marker genes, they identified eight different cell types, but several important and acknowledged niche populations (i. a., vascular cells) were not reported (Doetsch et al., 1997). Further studies focused on discrete cell types, for example, microglia (Ribeiro Xavier et al., 2015) and endothelial cells/vasculature, (Shen et al., 2004, 2008). But, to the best of our knowledge no publication as yet characterizes all SVZ cells together and gives an estimate on the proportions of cell types. The size of a given cell population depends on the choice of marker genes used for identification. For example, qNSCs are identified and isolated by co-expression of GFAP and PROM1, but not EGFR (Fischer et al., 2011). However, there is evidence for PROM1 negative qNSCs (Codega et al., 2014; Obernier et al., 2018), which in turn are difficult to distinguish from niche astrocytes. Here, instead of classifying cells based on few selected marker genes, we used genome wide expression profiling to cluster cells by transcriptome similarity and uncover cell identities by analyzing a multitude of genes. As the SVZ is not an enclosed tissue, it is challenging to obtain absolute numbers of the cellular composition. Nevertheless, our study provides an up to date characterization of cell types residing in the adult SVZ neurogenic niche including their gene expression signatures and a marker-independent estimate on the abundance of cell populations (Figures 6 and S1B). A user-friendly online tool makes our data easy accessible and allows to browse genes of interest in the context of 22 cell types.

Consistent with recent single-cell RNA sequencing studies, we resolved different NSC activation states (Basak et al., 2018; Dulken et al., 2017; Llorens-Bobadilla et al., 2015). In addition, our data allowed us to distinguish between niche astrocytes and NSCs. The cell type enriched genes identified in this study expand the existing pool of stem cell markers and help to discriminate different NSCs states from each other as well as from the other niche populations. RNA velocity analysis revealed that astrocytes and qNSCs have rather stable transcriptomes, whereas aNSCs are about to change their gene expression signature and move uniformly towards the TAPs state. mTAPs underwent vigorous transcriptomic changes probably due to their actively dividing state. Movements decreased from early to late NBs and made us hypothesize that progenitors in the SVZ have a less dynamic transcriptome once they acquired the NB fate. It is likely that RNA dynamics increase again when NBs reach the OB, where they terminally differentiate into interneurons.

**Figure 6.**
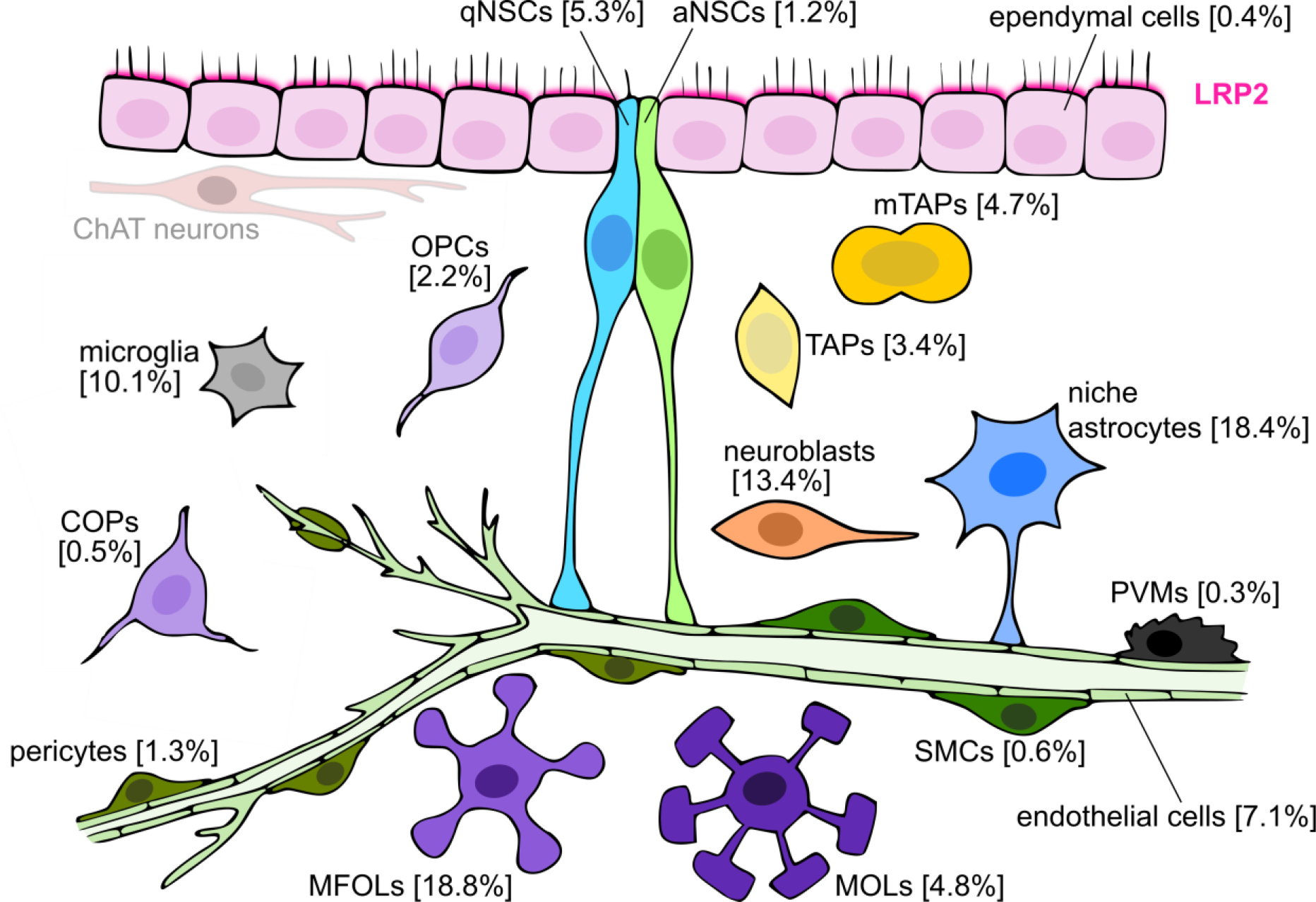
Cellular composition of the adult SVZ neurogenic niche and expression of LRP2. Illustration of cell types residing in the SVZ. Cells depicted with black outline were characterized in this study. Numbers in brackets are the mean of obtained cell type proportions. ChAT neurons (gray outline) could not be identified unambiguously in our data. LRP2, marked in red, is specifically expressed in ependymal cells (Gajera et al., 2010).

**Abbreviations:** qNSCs: quiescent neural stem cells; aNSCs: activated neural stem cells; TAPs: transient amplifying progenitors; mTAPs: mitotic TAPs; PVMs: perivascular macrophages; SMCs: smooth muscle cells; OPCs: oligodendrocyte progenitor cells; COPs: differentiation-committed oligodendrocyte precursors; MFOLs: myelin forming oligodendrocytes; MOLs: mature oligodendrocytes.

NSCs have been postulated to be tri-potent and able to generate neurons, oligodendrocytes, and astrocytes. This assumption has been largely based on *in vitro* experiments, which demonstrated that adult NSCs are capable to self-renew and to produce different cell types (Pastrana et al., 2011; Reynolds and Weiss, 1992). *In vivo* population fate-mapping studies showed that SVZ NSCs give rise to OPCs and mature oligodendrocytes (Menn et al., 2006), but clonal analysis revealed only the production of interneurons from individual NSCs (Calzolari et al., 2015). Along this line, the sequencing data, which reflects a snapshot of processes taking place *in vivo*, revealed the neurogenic lineage, but no connection between NSCs and OPCs. However, as oligodendrogenesis is a rare event (Menn et al., 2006), it is conceivable that, despite the vast cell type coverage and high resolution of our dataset, the sampling was not dense enough to capture intermediate cells that support this process. As single-cell RNA sequencing methods and computational tools are developing rapidly (reviewed in Kester and van Oudenaarden, 2018), *in silico* lineage tracing of adult NSCs will most likely improve.

Our unbiased approach makes it possible to study the SVZ in basically all adult neurogenesis mouse models including conditions of disturbed neurogenesis. Here, we studied the consequences of LRP2-deficiency on the adult SVZ neurogenic niche. Single-cell transcriptomics reproduced the main findings of a previous immunohistological study (Gajera et al., 2010) and revealed unexpected novel insights to the function of LRP2 in controlling adult neurogenesis. In mice lacking LRP2, the number of proliferating cells is reduced and BMP4 signaling is increased (Gajera et al., 2010). We expanded these findings by showing that TAPs expressed less cell cycle genes and that *Id3*, a downstream target of BMP4, was increased in NBs of KO mice. In addition, we also detected a global decrease in expression of ribosomal proteins in cells derived from LRP2-deficient SVZ, which was most pronounced in TAPs, and a reduction in Wnt activity. LRP2 is known to modulate signaling by BMP4 and SHH by acting as a retrieval receptor for these morphogens from the extracellular space (Christ et al., 2012, 2015; Gajera et al., 2010). A possible involvement of this receptor in Wnt signaling in the SVZ identified by single-cell RNA transcriptomics is novel and exciting, and supports a model whereby LRP2 acts as a versatile clearance receptor in ependymal cells that binds a variety of morphogens and regulates their availability in the SVZ stem cell niche. Upon LRP2-deficiency, the morphogen levels are out of balance leading to dysfunctional signaling and perturbed neurogenesis in the SVZ.

## ACKNOWLEDGMENTS

We thank Susanne A. Wolf for her help with animal experiments and optimizing tissue dissociation, Christine Kocks for guidance and support with single-cell transcriptome profiling, Anastasiya Boltengagen, Salah Ayoub, and Maria Schmeisser for technical support, Sten Linnarsson and Gioele La Manno for their help with the velocyto.py package, and Dominic Grün and F. Alexander Wolf for testing their methods on our data. We thank Christine Kocks and Monika Piwecka for critically reading of the manuscript and the entire Rajewsky lab for critical discussions, helpful comments, and sharing code. A.M. acknowledges funding from the European Union’s Horizon 2020 research and innovation programme, under the Marie Sklodowska-Curie Actions (MSCA) grant agreement No 721890.

## AUTHOR CONTRIBUTIONS

V.Z. initiated the study, performed experiments, analyzed data, and prepared the manuscript with the help of N.R. and A.M.. A.M. performed the computational analyses assisted by V.Z.. L.B. performed and evaluated immunohistochemistry experiments. T.W. and N.R. supervised the study. N.R. was responsible for the conceptual design and study coordination. All authors were involved in data interpretation.

## DATA AND SOFTWARE AVAILABILTY

Data have been deposited in GEO under accession number GEO: GSE111527.

## SUPPLEMENTAL INFORMATION

Supplemental Information includes seven figures and four tables.

## EXPERIMENTAL PROCEDURES

### Animals

Experiments involving animals were performed according to institutional guidelines following approval by local authorities (X9017/12). Mice were housed in a 12 hour light-dark cycle with *ad libitum* food and water. We used adult mice (2-4 month of age) for our experiments. For Dataset A of the Drop-seq experiments, we used female and male C57BL6/N mice (overview in Figure S1A). For Dataset B, we used male LRP2-deficient mice (KO) and matched littermate controls (Ctr) (overview in Figure S5A). KO mice were compound heterozygous for two different *LRP2 null* alleles. One allele (LRP2^+/-^) was derived by targeted gene disruption (Spoelgen et al., 2005). This line was crossed with the *Tcf/Lef_LacZ* reporter strain (Mohamed et al., 2004). The second mutant allele (Lrp2^267/+^) was derived in an ENU mutagenesis screen (Zarbalis et al., 2004). LRP2 KO animals *(Tcf/Lef_LacZ, Lrp2^267/-^)* were derived from F1 crosses of (*Tcf/Lef_LacZ, Lrp^+/^-*) and *Lrp2^267/+^* animals. As no obvious phenotypes were observed in mice carrying only one targeted *Lrp2* allele, both heterozygous as well as wildtype animals were used as controls (referred to as Ctr throughout the study).

### Generation of SVZ single-cell suspension

For each Drop-seq experiment, three to five mice were deeply anesthetized by intraperitoneal injection of pentobarbital (100 µl of 160 mg/ml pentobarbital sodium solution per mouse) and subsequently transcardially perfused with 20-25 ml ice cold NaCl/Heparin solution (40 µl Heparin-Natrium (250,000 I. E./10 ml) in 500 ml 0.9% NaCl) until the liver was pale. Brains were immediately extracted and collected in ice cold Hanks´ Balanced Salt Solution (HBSS) without Ca^2+^ and Mg^2+^ (Sigma-Aldrich #55021C). The SVZ of the lateral ventricles was microdissected as described in (Walker and Kempermann, 2014). SVZ from up to five brains was pooled for one enzymatic dissociation reaction using the Neural Tissue Dissociation Kit (P) (Miltenyi #130-092-628) with minor changes: for enzyme mix 1, 1900 µl buffer X was supplemented with 0.96 µl 0.143 M beta-mercaptoethanol (final concentration 70 µM) and 7.8 µl 10% bovine serum albumin (BSA; final concentration 0.04%; molecular biology grade) one day in advance and stored at 4°C. 50 µl enzyme P was added briefly before use and the enzyme mix was pre-warmed to 37°C. SVZ tissue was slightly minced on ice and transferred into a 15 ml tube using the pre-warmed enzyme mix and a 1000P pipette with the tip cut in front. After 15 minutes incubation at 37°C (slowly rotating), enzyme mix 2 was added and the tissue carefully dissociated by pipetting ten times up and down with a 1000P pipette. The suspension was incubated another 10 minutes at 37°C (slowly rotating) and dissociated again by careful pipetting as before. In cases of remaining tissue clumps, a 200P pipette tip was placed in front of the 1000P tip and few additional, very careful pipetting steps were added. Subsequently, the enzymatic digestion was stopped by adding 5 ml room temperature HHB solution (HBSS with Ca^2+^ and Mg^2+^ (Sigma-Aldrich #55037C), 10 mM HEPES, 0.01% BSA). Cells were filtered through a 70 µm cell strainer (Sigma #CLS431751-50EA; pre-wet with 1 ml HHB). The previous cell tube and the filter were washed twice with 1 ml HHB. After centrifugation for 10 minutes at 10°C and 350xg, cells were resuspended in HHB for the following 3-density centrifugation step to remove small cellular debris (based on Guez-Barber et al., 2012), in the following referred to as percoll gradient). If tissue from four to five mice was used, three percoll gradients (in three 15 ml tubes) were processed in parallel and cells were resuspended in 3 ml HHB (tissue from 2-3 mice was resuspended in 2 ml HHB for two individual percoll gradients). To form the 3-density gradient, 1 ml of each ice cold solution (high density: 19% percoll (Sigma #P1644), 22.5 mM NaCl in HBSS/HEPES (HBSS with Ca^2+^ and Mg^2+^, 10 mM HEPES); medium density: 15% percoll, 17.7 mM NaCl in HBSS/HEPES; low density: 11% percoll, 13.8 mM NaCl) was carefully layered on top of each other in a 15 ml Falcon tube on ice starting with the high density solution at the bottom. Last, 1 ml of the filtered cell suspension was applied on top. After 3 minutes centrifugation at 430xg and 4°C, the cloudy top layer containing debris and dead cells was removed and discarded (about 2 ml per tube). Subsequently, cells were pelleted by 5 minutes centrifugation at 550xg and 4°C, resuspended in ice cold PBS/0.01% BSA, pooled and filtered through a pre-wet 20 or 30 µm strainer (20 µm: HiSS Diagnostics #43-50020-03, 30 µm: Miltenyi #130-041-407). Remaining tubes and strainer were washed twice with 0.1 ml PBS/0.01% BSA. Finally, cells were counted using trypan blue (Sigma # T8154) in a Neubauer counting chamber and either diluted to 100 cells/µl using PBS/0.01% BSA (proceed with Drop-seq directly), or fixed in methanol as described below for later use.

### Fixation and rehydration of cells for Drop-seq

In two out of nine Drop-seq experiments, living, single-cells were fixed in methanol and therefore preserved for later experiments (according to Alles et al., 2017). In short, the final single-cell suspension was centrifuged for 5 minutes at 300xg and 4°C to reduce the volume to 200 µl. The single-cell suspension was vortexed gently and 800 µl 100%, −20°C cold methanol was added dropwise to reach a final methanol concentration of 80%. After 15 minutes incubation on ice, the cell suspension was transferred to a 1.5 ml Eppendorf tube, sealed with Parafilm and stored at −20°C or −80°C until use. For Drop-seq, cells were equilibrated on ice for 5-10 minutes and subsequently centrifuged for 10 minutes at 500xg and 4°C. The supernatant was discarded, and cells resuspended in 0.5 ml rehydration solution (PBS, 0.01% BSA, 1:40 RiboLock; Fisher Scientific #EO0382). Cells were filtered through a 30 µm filter and diluted to either 50 cells/µl (experiment an003_F) or 100 cells/µl (experiment an002) with rehydration solution. The cell recovery rate was about 50%.

### Drop-seq procedure, single-cell library generation and sequencing

Drop-seq was performed as described in Alles et al., 2017 (based on Macosko et al., 2015). In brief, monodisperse droplets with a volume of about 1 nl were generated using a self-built Drop-seq device. After SMART PCR, cDNA libraries had an average size of 1.2 to 2.0 kb. Final single-cell Drop-seq libraries had an average size of 600 to 780 bp and were sequenced at a concentration of 1.8 mM on Illumina NextSeq500 sequencers with 1% PhiX spike in for run quality control. We used Illumina Nextseq500/550 High Output v2 kits (75 cycles) and sequenced in paired-end mode: Read 1: 20 bp (bases 1-12 cell barcode, bases 13-20 UMI); Read 2: 64 bp; Index read: 8 bp.

### Convention for experiment IDs

Nine Drop-seq samples were produced in this study. The experiment IDs (exp_ID) for Dataset A are an002, an003_F, an003_L, an008 and an009. For Dataset B, the samples were referenced as an007_WT, an007_KO, an010_WT, an010_KO. The samples an007_KO and an010_KO were derived from LRP2-deficient mice (KO) and an007_WT and an010_WT from wildtype and heterozygous littermate controls (Ctr), as described in the experimental procedures. an007_WT contained cells derived from one wildtype and three heterozygous mice. For an010_WT, two wildtype and two heterozygous mice were used.

### Drop-seq computational pipeline: data processing, alignment and gene quantification

Sequencing quality was assessed by FastQC. To produce the digital gene expression matrix (DGE), we used the Drop-seq tools v1.12 and Picard-tools v2.9.0, following the standard pipeline in the Drop-seq core computational protocol v1.2 with default parameters (based on Macosko et al., 2015). In brief: We extracted the UMIs and cell barcodes from read 1 while discarding it, added them as metadata information (tags) to the corresponding sequence of read 2, and performed the downstream analysis as in single-end mode. Reads with low quality barcodes were discarded. We trimmed potential SMART adapter contaminants and poly(A) stretches. Subsequently, reads were aligned to the mouse genome mm10 using STAR v2.5.3a with default parameters and annotated according to GRC38m.p4. Typically, 70-90% of reads mapped to the genome. Non-uniquely mapped reads were discarded. We corrected reads for bead synthesis errors such as missing last base of the cell barcode. For each sample, the number of cells was estimated by plotting the cumulative distribution of reads per cell against the barcodes sorted by descending number of reads and calculating the inflection point (“knee”). The inflection point was calculated considering the top 25,000 barcodes for all samples except an007_KO and an007_WT for which the top 15,000 barcodes were considered. Finally, we extracted the DGE for each sample.

### Cell and gene filtering

For downstream analysis, we excluded cells which expressed less than 500 UMIs, less than 200 genes, and had more than 10% of total UMIs quantified from mitochondrial genes. Furthermore, we removed genes which were expressed in less than three cells. All samples were analyzed as described in the next paragraph. After individual analysis and comparison of replicates, DGEs from replicates were combined. For Dataset A, the DGEs of five independent Drop-seq runs were pooled. We excluded 91 cells, which formed four small cell clusters consisting of one to three replicates, only. After filtering, Dataset A comprised 9,804 cells, which were analyzed further. In Dataset B, the DGEs of 4 Drop-seq runs (two replicates with LRP2-knockout and littermate control derived cells processed in parallel) were pooled resulting in 8,056 cells for further analysis.

### Normalization, clustering, marker discovery, and data visualization

The following analysis was performed using Seurat v1.4. To normalize UMI counts for every gene per cell, we divided its UMI counts by the total number of UMIs in that cell, multiplied the value by 10,000 and applied a logarithmic transformation. Using the MeanVarPlot function, we selected the genes that showed most variation across cells, with an average expression between 0.01 and 3 normalized UMI counts, and a minimum standard deviation of 1. With these genes we performed Principal Component Analysis (PCA) to further reduce the dimensions of the data. By applying the jackstraw function, we identified the significant first principal components, which were used for further analysis. We used this PCA transformation to perform clustering with SNN-cliq. We evaluated which resolution parameter is appropriate by testing several different parameters in combination with the AssesNode function. The resolution parameter determines in how much detail cells are partitioned into clusters, whereas the AssessNode function merges clusters, which are not meaningful. In the end, Dataset A was clustered with a resolution of 1.0 and Dataset B with 0.8, respectively. Dendrograms representing cluster relationships were produced with the BuildClusterTree function. For data visualization, we performed t-distributed stochastic neighbor embedding (tSNE). To identify cell type enriched genes, we applied the FindAllMarkers function with the parameters min.pct and thresh.use set to 0.25. To identify significantly up- and downregulated genes in neural stem cells compared to astrocytes, we used the FindMarkers function. The results were visualized in heatmaps produced with the DoHeatmap function. The expression of single genes was depicted using custom R scripts, either per cell cluster as distribution of normalized UMI counts (violins) or per cell as color gradient in tSNE space. For better visualization, we added noise to the normalized UMI counts in Figure S4F.

### RNA velocity analysis

To calculate the RNA velocity, we applied the velocyto python package (La Manno et al., 2017). Velocyto counts the spliced and unspliced reads separately. After normalization, variable gene selection, and smoothing/imputation, the method uses all cells to estimate the expected steady state ratio between spliced and unspliced molecules. From here, velocyto calculates and assigns an RNA velocity value for each cell per gene to extrapolate the future transcriptional cell state.

We run the Command Line Interface (CLI) of velocyto (version 0.11.0) in permissive mode. Using all cells from Dataset A, we normalized, selected the top 1000 variable genes further thresholding for minimum expression, performed data imputation with a neighborhood of 200 cells, and calculated the velocities. All steps were performed following the built-in functions. We then isolated the cells of the neurogenic lineage based on the clustering by Seurat (as described above) and plotted their velocities. Finally, we estimated the differentiation starting point of the selected cells by using the backward Markov process on the transition probability matrix to determine high density regions. All steps were performed with default parameters.

### Subclustering

To produce the subclusterings of Dataset A and Dataset B, cells belonging to the clusters of interest were isolated from the DGE and the Seurat analysis was repeated with the clustering resolution set to 0.8 and 0.7, respectively.

### Gene set scoring

For cell type characterization and comparison of our data with the literature, we downloaded previously published gene sets (Llorens-Bobadilla et al., 2015) and scored our cells as follows: Genes, which were not in the gene sets or had an expression of less than 50 UMIs across all cells of interest were discarded from the normalized DGE. The expression range of remaining genes was linearly transformed to (0, 1) range for every gene. The score per cell was calculated by averaging the transformed gene expression values of all genes from a certain gene set. The distribution of scored cells was plotted as violin per subcluster and gene set.

### Proliferation analysis of LRP2-deficient vs. control cells (Dataset B)

Following the method above (gene set scoring), we scored all cells from Dataset B according to cell cycle using a published gene set (Kowalczyk et al., 2015). Per cell cluster and condition (LRP2 KO vs. Ctr), we calculated the cumulative fraction of cells against the score. We performed Kolmogorov-Smirnov test comparing the cumulative distributions of the two conditions for each cluster to determine statistical significance of data.

### Differential gene expression analysis

To identify genes, which are deregulated in cells derived from LRP2-deficient SVZ as compared to control cells, we performed differential gene expression analysis (DE) in each cell cluster using EdgeR (Robinson et al., 2010). We analyzed both replicates independently (an007_WT vs an007_KO and an010_WT vs an010_KO). Each cell was considered as a sample. We compared the gene expression of cells belonging to the same cluster between conditions (knockout vs. control). As normalizing factor, we used the total UMIs of each cell. Here, we report only genes, which had a p-value smaller than 0.01 and a fold change into the same direction in both replicates (an007 and an010). Data was visualized using MA plots: y axis represents the log2 ratio between the averages of a genes expression in knockout vs. control derived cells (log2FC), x axis represents the log2 average UMI expression over control cells. To all averages a pseudocount of 1 was added.

### Ribosomal gene expression comparison in cells derived from LRP2-deficient SVZ vs. control cells

We compared the expression of ribosomal genes vs. the expression of remaining genes in cells derived from LRP-deficient vs. control SVZ. First, we discarded genes with an average expression lower than 0.5 log2 (mean normalized UMI counts) in control cells. Second, we separated ribosomal genes and all remaining genes. We calculate the log2FC, as described in the MA-plot paragraph. The distributions of the two gene sets were plotted separately for each cluster as violin plots. Statistical significance of data was determined using unpaired student’s t-test.

### Generalized linear mixed model

To test whether cluster proportions change due to genotype (LRP2-deficient or control), we employed a generalized linear mixed model (GLMM) with a binomial distribution. The GLMM can estimate the effects of each factor (e.g., condition or batch) on cell type proportions. We modeled the condition as a fixed effect and the batch as a random effect. We used a binomial distribution because the response for every cell is binary (meaning every cell either belongs to the given cluster or not). We employed a Laplace approximation to estimate the parameters of the model, using the glmer function of the lme4 package in R (Bates et al., 2015).

### BrdU labeling experiments

To label proliferating cells in the adult mouse brain, animals were injected intraperitoneal once with BrdU (50 mg/kg body weight) and sacrificed 24 hours later by intraperitoneal injection of pentobarbital (as before) and subsequent transcardial perfusion with 4% paraformaldehyd. Brains were collected and processed for routine paraffin embedding and sectioning. 10 µm sections were washed with 2.4% H_2_O_2_ in 1xTBS for 30 minutes, treated with 2 N HCl at 45°C for 1.5 hours, and neutralized by washing in 0.1 M borate buffer (pH 8.5) for 15 minutes at room temperature. Sections were blocked in 10% donkey serum in 1xTBS/0.3% Triton X-100 for 1 hour, and subsequently incubated with rat anti-BrdU antibody (1:500; AbD Serotec) overnight at 4°C, followed by donkey anti-rat Biotin SP antibody (1:250; Jackson Immuno Research) for 2 hours, and finally 1 hour in ABC-Elite Reagent (PK-6100; Vector Labs). The color reaction was performed using Ni-diamino benzidine.

### Immunohistology

Standard immunohistology was performed on free-floating sections. Therefore, PFA fixed brains were infiltrated with 30% sucrose in PBS for 48 hours, cut into 40 µm sections, and stored in cryoprotectant (25% Glycerol, 25% Ethylene Glycol, 50% Phosphate Buffer pH 7.8) at −20°C until further use. Tissue sections were incubated with primary antibodies at the following dilutions: rabbit anti-S6-RP (1:50; Cell Signaling) or rabbit anti-ID3 (1:100; Abcam). Bound primary antibody was visualized using secondary antisera conjugated with Alexa-488 (1:500; Invitrogen) for S6-RP or the tyramide signal amplification kit (PerkinElmer) for ID3. Pictures were taken on a confocal SPE microscope. We counted individual ID3 positive cells and quantified the mean gray value for S6-RP in the SVZ using the software ImageJ.

### LacZ staining

For LacZ staining, dissected brains were fixed for 3 hours in 4% PFA and infiltrated with 30% sucrose in PBS for 48 hours. Tissues were embedded in Tissue-Tek OCT (Sakura, Japan), cooled down on dry ice, sectioned at 12 μm thickness on a rotary cryotome (Leica, Germany), and stored at −20°C until further use. To start the staining procedure, slides were thawed for 5 minutes at room temperature and sections subsequently fixed by incubating in 1xPBS (pH 7.4), 2 mM MgCl_2_, 5 mM EGTA, 0.2% Glutaraldehyd for 5 minutes. Following washing steps with 1xPBS plus 2 mM MgCl_2_, sections were permeabilized with 1xPBS, 0.02% NP40, 0.01% Sodium Deoxycholate, 2 mM MgCl_2_ for 10 minutes, and incubated overnight at 37°C in 1 mg/ml X-GAL (5-Bromo-4Chloro-3-indolyl-B-D-galactopyranoside) solved in staining solution (1xPBS, 20 mM Tris (pH 7.3), 0.02% NP40, 0.01% Sodium Deoxycholate, 2 mM MgCl2, 5 mM Potassium Ferrocyanide). Images were acquired on a bright field microscope (Olympus BX51TF).

## SUPPLEMENTARY FIGURES

**Figure S1.**
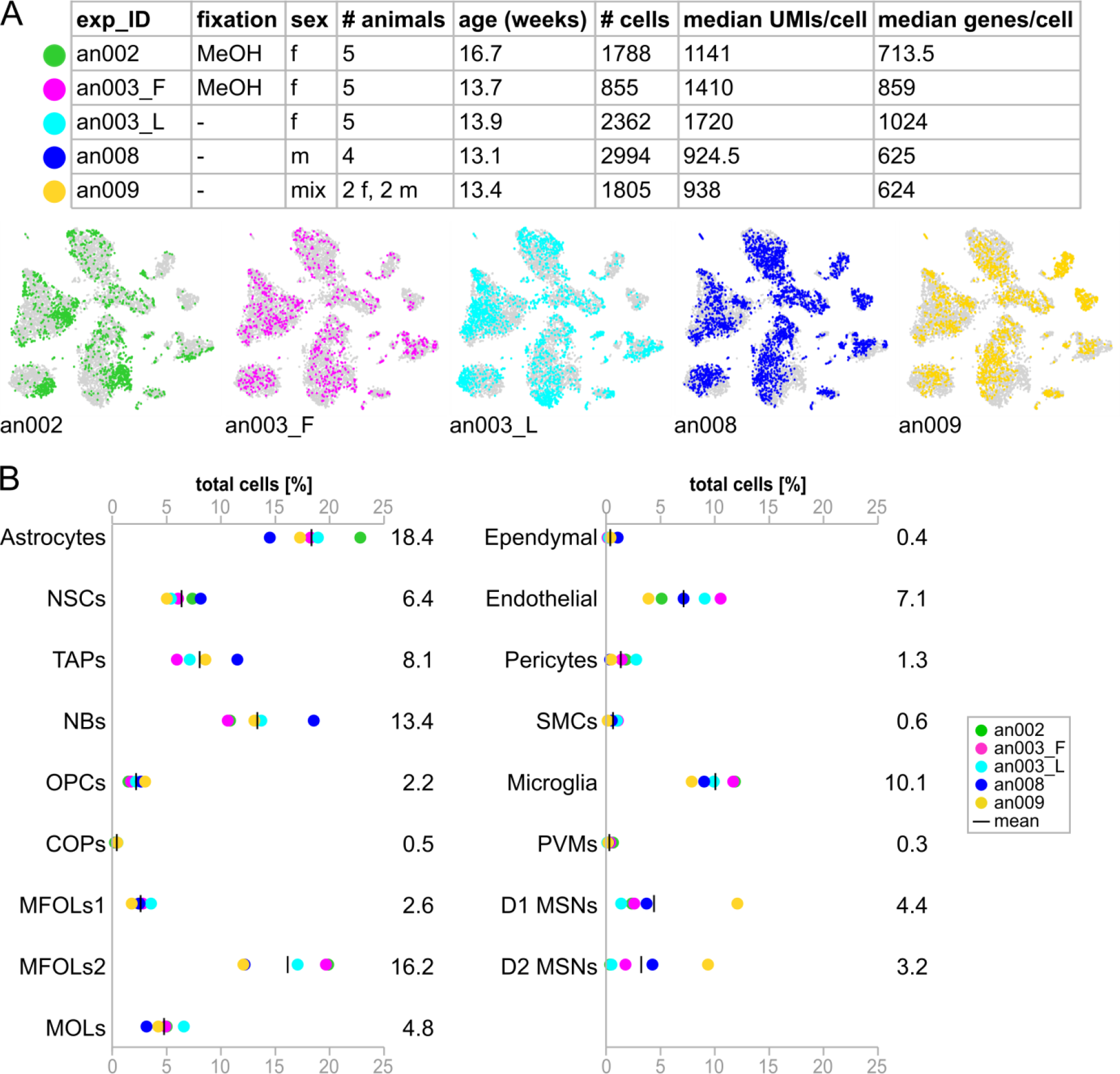
Cluster composition and distribution of replicates across cell types. Related to Figure 2. (A) Overview of replicates and distribution of replicates in the all cell tSNE plot. Cells from one replicate are colored and plotted on top of all other cells (in gray). (B) Proportions of cell types. Numbers represent the mean.

**Abbreviations:** exp_ID: experiment ID; f: female; m: male; UMIs: unique molecular identifiers. Cell type abbreviations as in Figure 2.

**Figure S2.**
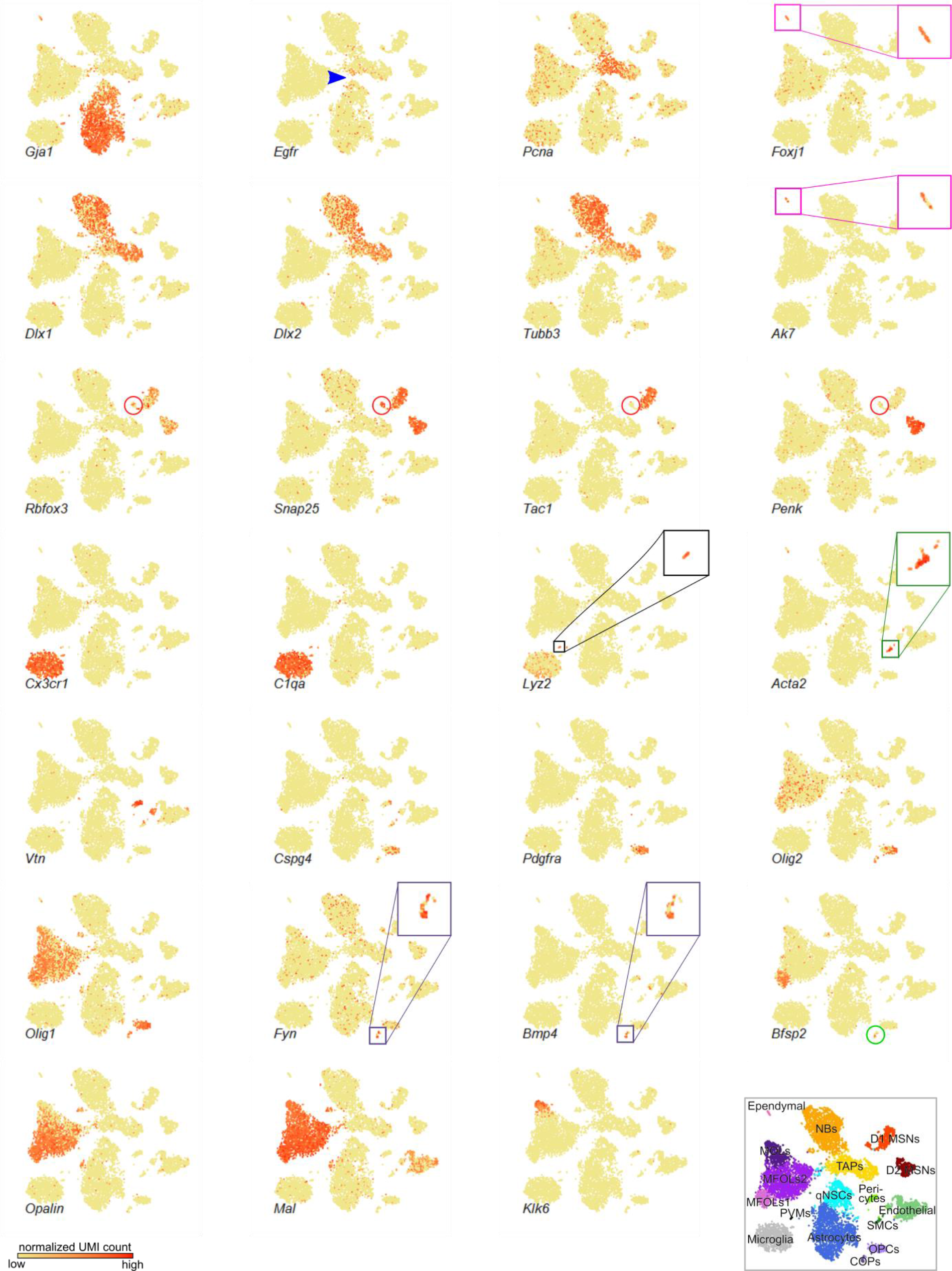
Marker gene expression. Related to Figure 2. tSNE plots of 9,804 cells colored by the expression of certain marker genes (yellow indicated low, red indicates high expression): *Gja1*: astrocytes and NSCs; *Egfr* (blue arrow) activated NSCs and neural progeny; *Pcna*: proliferating cells; *Foxj1, Ak7*: ependymal cells; *Dlx1, Dlx2*: committed neural progenitors; *Tubb3*: neuroblasts; *Rbfox3, Snap25*: mature neurons; *Tac1*: D1 MSNs; *Penk*: D2 MSNs; *Cx3cr1*: microglia; *C1qa*: immune cells (microglia and PVMs); *Lyz2*: PVMs; *Acta2*: SMCs; *Vtn*: pericytes; *Cspg4*: pericytes and OPCs; *Pdgfra, Olig2*: OPCs; *Olig1*: oligodendrocytes; *Fyn, Bmp4*: COPs; *Opalin, Mal*: MFOLs; *Klk6*: MOLs. *Bfsp2* is a putative new marker gene to discriminate MFOLs1 from MFOLs2. *Bfsp2* was highest expressed in MFOLs1, but also detected in COPs (green circle), indicating that MFOLs1 represent an earlier stage of myelin forming oligodendrocytes than MFOLs2. Classification of oligodendrocyte clusters was based mainly on genes published in (Marques et al., 2016). Squares are close ups of certain cell clusters. Red circles enclose putative ChAT neurons. At the bottom right: tSNE colored by cluster annotation. Cell type abbreviations as in Figure2.

**Figure S3.**
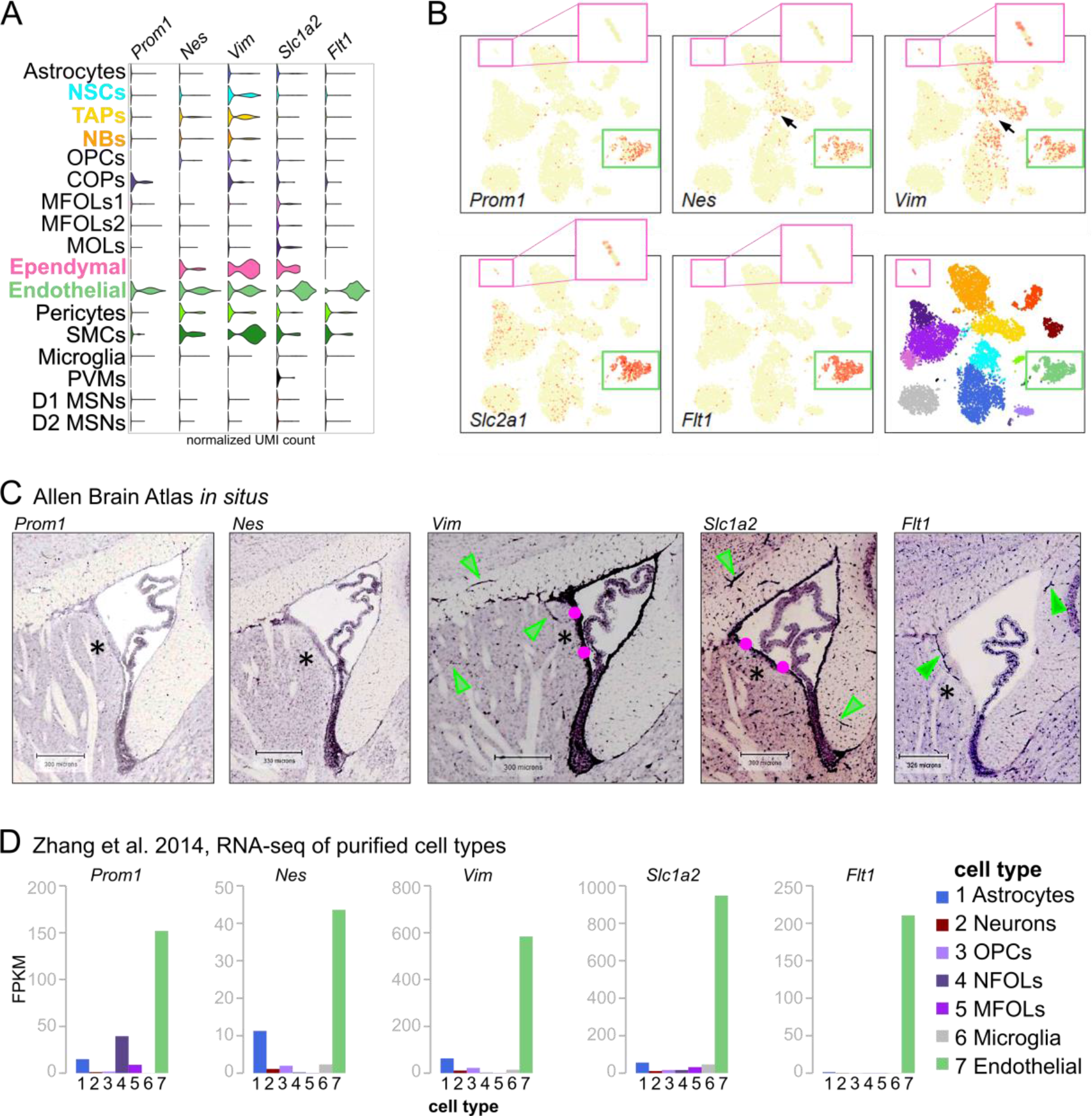
Characterization of endothelial cells. Related to Figure 2. (A, B) Visualization of genes associated with ependymal, endothelial, neural stem and progenitor cells. (B) Pink boxes highlight and zoom into the ependymal cell cluster. Endothelial cells are highlighted by green boxes. Arrows point at neural progenitors. (C) *In situ* images from the Allen Brain Atlas (Lein et al., 2007) depicting the expression of genes from A and B in the adult mouse brain. *Prom1* and *Nes* mRNA was barely detected. Green arrowheads point to blood vessels. Pink circles highlight staining of the ependymal cell layer. Asterisks label the lateral wall. (D) Expression of genes from A-C in published bulk RNA-sequencing data from sorted cell populations (data from Zhang et al., 2014). Abbreviations as in Figure 2.

**Figure S4.**
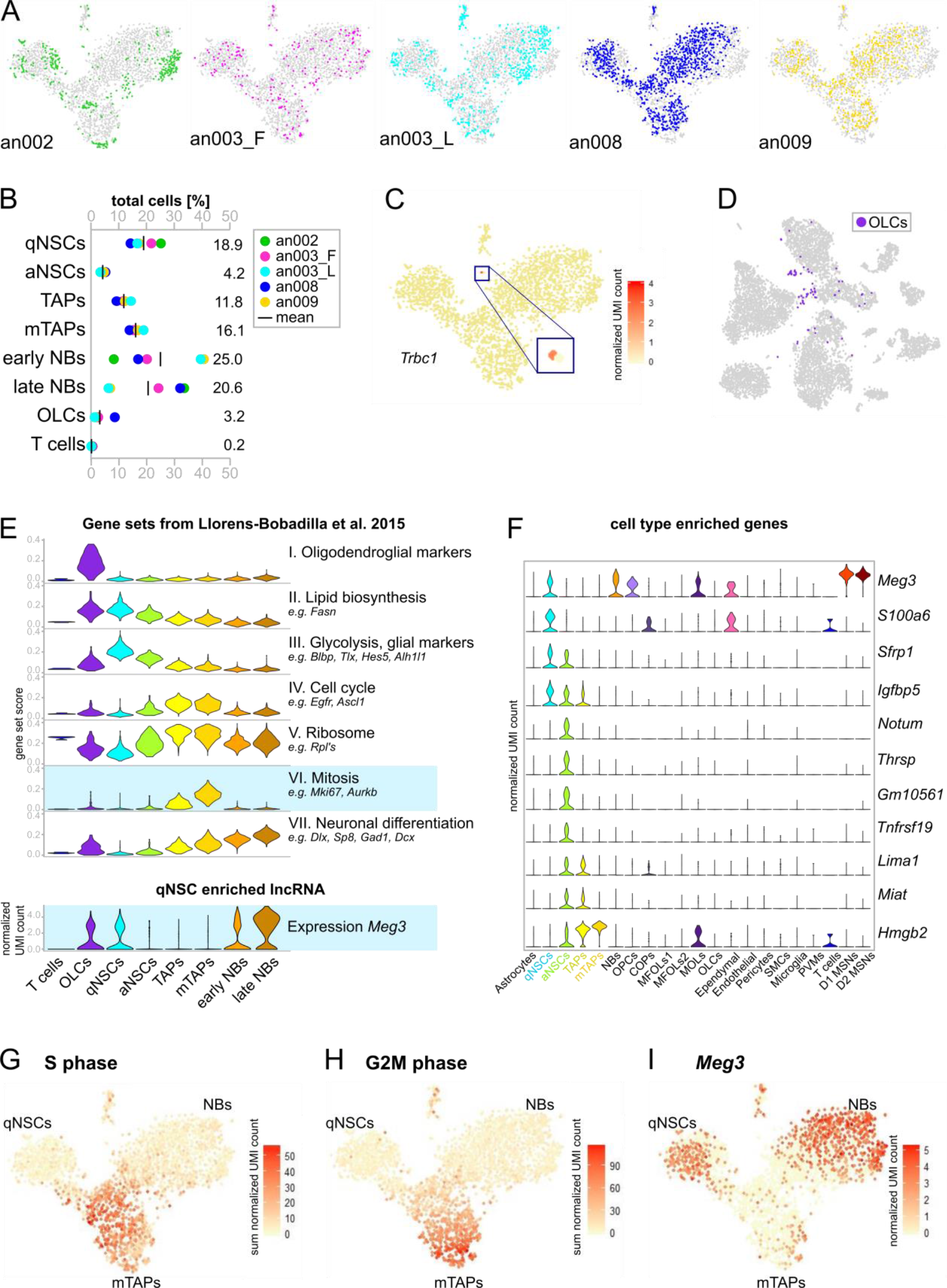
Subcluster characterization and NSC activation stage enriched genes. Related to Figure 3. (A) Distribution of replicates in the tSNE plot of the subclustering. Cells from one replicate are colored and plotted on top of all other cells (in gray). (B) Proportions of cell types after subclustering. Numbers represent the mean. (C) tSNE plot of the subclustering colored by the expression of T cell receptor *Trbc1*. The expression of T cell receptors was specific for subcluster 1 (blue square represents a close up) disclosing these cells as contaminating T cells. Of note, the subcluster consists of four cells deriving from four different replicates. (D) Localization of oligodendrocyte-like cells (OLCs, in purple) in the all cell tSNE. In the subclustering analysis, subcluster 2 separated due to oligodendrocyte associated genes (gene set I, panel E). However, OLCs also expressed some glial and neuroblast genes (gene sets II, III and VII, panel E) and localized close to NSCs, TAPs, and NBs in the all cell tSNE (panel D) suggesting that OLCs might represent cell doublets. But, two previous single-cell RNA sequencing studies also observed OLCs within their FACS-sorted populations (Dulken et al., 2017; Llorens-Bobadilla et al., 2015). Therefore, we doubt that these cells are doublets, but hypothesize that they might represent a so far uncharacterized cell type. (E) Characterization of subclusters based on gene sets published in (Llorens-Bobadilla et al., 2015). At the bottom: the expression of the long non coding RNA *Meg3* negatively correlates with the expression of mitotic genes (blue background). See also panels G-I. (F) NSC activation stage enriched genes in the context of all identified cell types from Figures 2A and 3C. (G, H) Expression sum of S (G) and G2M (H) phase associated genes (from Kowalczyk et al., 2015) in the tSNE plot of the subclustering. (I) Expression of the long non coding RNA *Meg3* in the tSNE plot of the subclustering.

**Abbreviations:** qNSCs: quiescent neural stem cells; aNSCs: activated NSCs; TAPs: transient amplifying progenitors; mTAPs: mitotic TAPs; NBs: neuroblasts; OLCs: oligodendrocyte-like cells. Other cell type abbreviations as in Figure 2.

**Figure S5.**
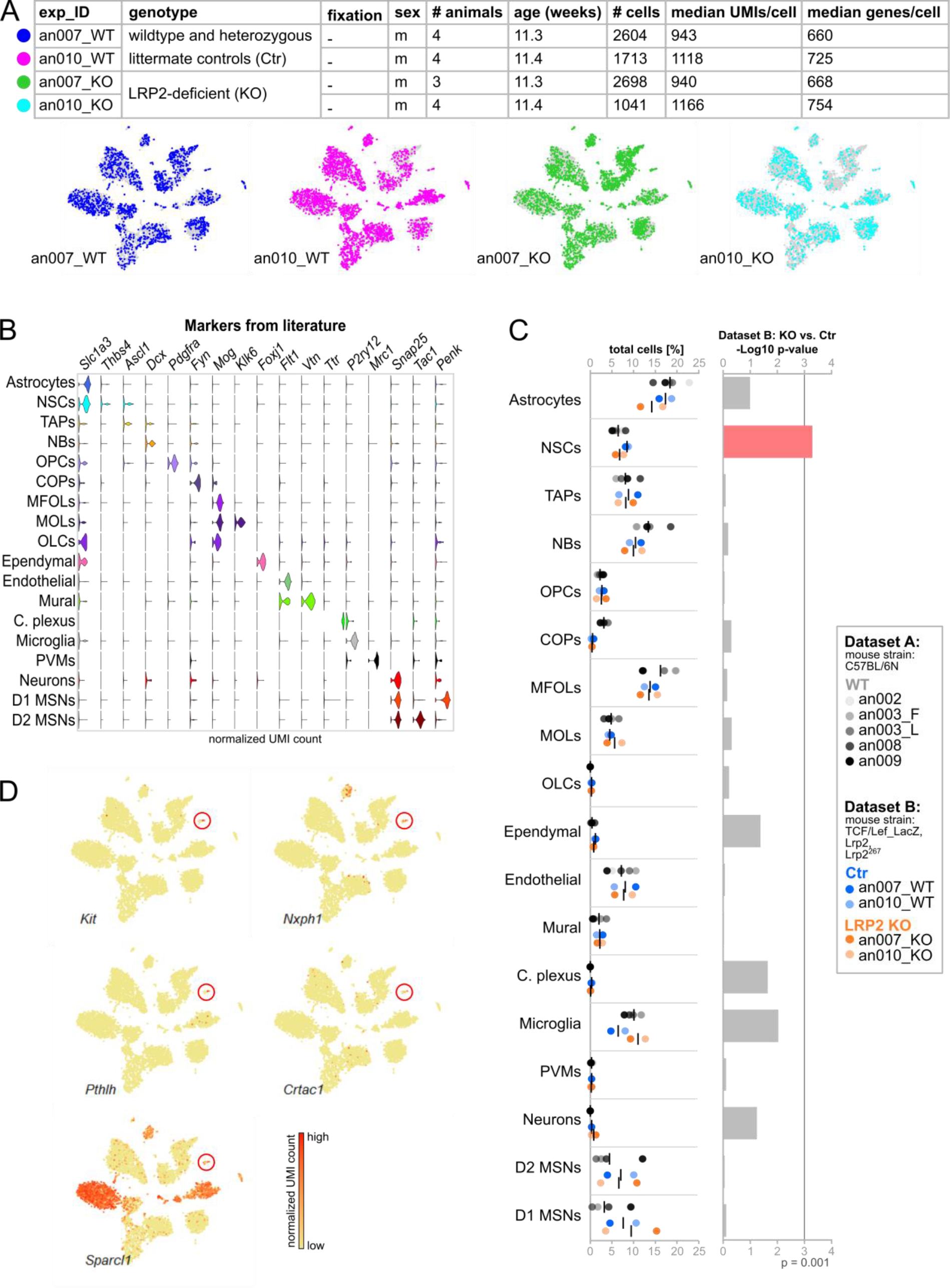
Cluster composition, cell type identification and distribution of replicates across cell types in the LRP2 KO/Ctr Drop-seq analysis (Dataset B). Related to Figure 4. (A) Overview of replicates and distribution of replicates in the all cell tSNE plot. Cells from one replicate are colored and plotted on top of all other cells (in gray). an007_WT contained cells derived from one wildtype and three heterozygous mice. For an010_WT, two wildtype and two heterozygous mice were used. (B) Identification of cell types based on known marker genes. (C) Proportions of cell types in Dataset A (gray, wildtype) and Dataset B (blue: control, orange: LRP2-deficient mice). Numbers represent the mean. Right panel: P-values from the comparison of cell type proportions in Dataset B: Ctr vs. KO. Statistical significance of data was determined using a generalized linear mixed model. The p-value for NSCs is highlighted in red.

**Abbreviations:** exp_ID: experiment ID; m = male; KO: LRP2-deficient mice, Ctr: wildtype and heterozygous littermate controls; C. plexus: choroid plexus, OLCs: oligodendrocyte-like cells, other cell type abbreviations as in Figure 2.

**Figure S6.**
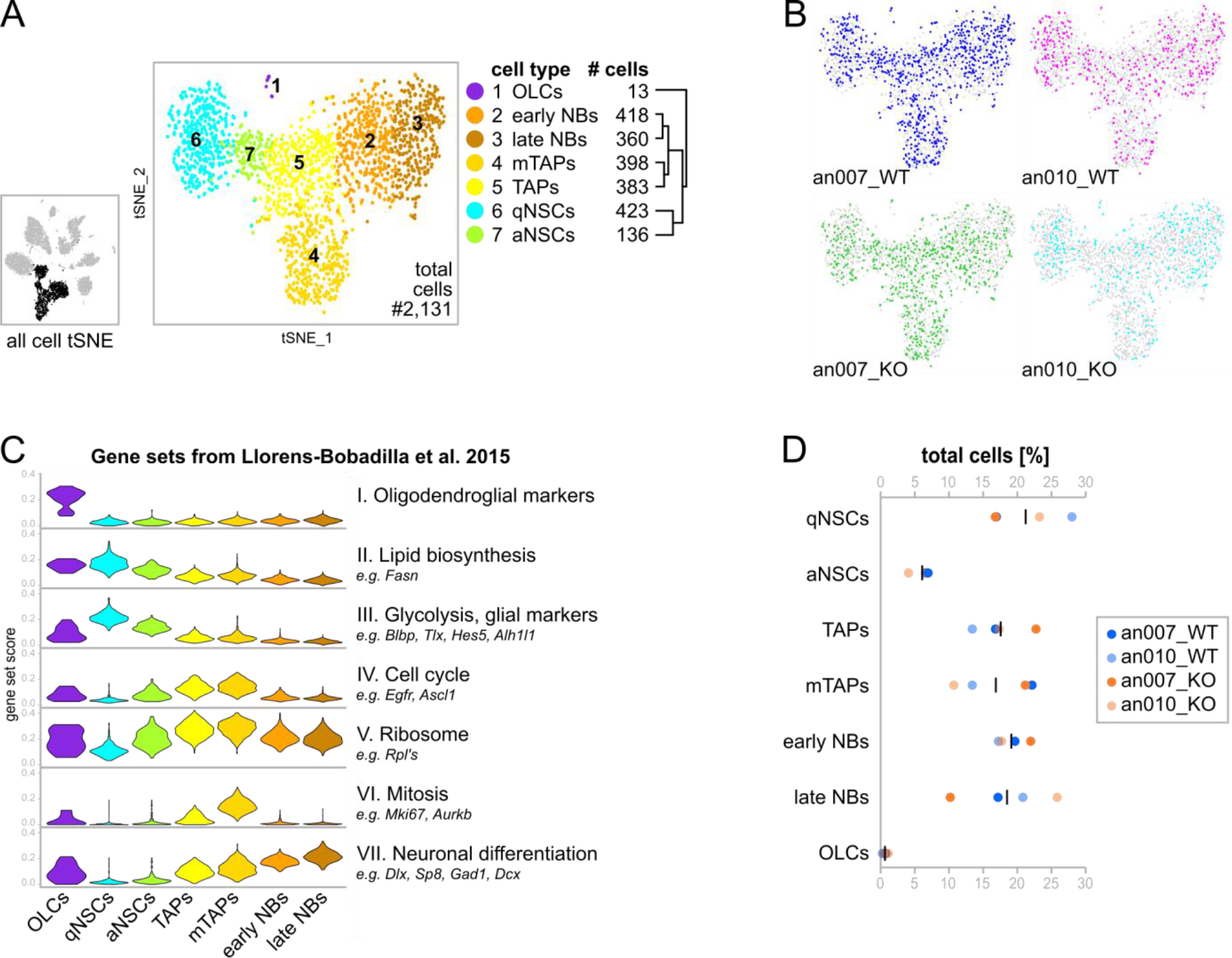
Subclustering of LRP2 KO/Ctr Data (Dataset B). Related to Figure 4. (A) tSNE plot of the subclustering analysis of 2,131 cells (NSCs, TAPs, and NBs in the all cell tSNE, on the left in black) reveals 7 cell clusters. Clusters 2-7 belong to the neurogenic lineage. The dendogram shows the relationship of cell types. (B) Distribution of replicates in the tSNE plot of the subclustering. Cells from one replicate are colored and plotted on top of all other cells (in gray). (C) Cell type characterization based on gene sets published in (Llorens-Bobadilla et al., 2015). (D) Proportions of cell types after subclustering. Numbers represent the mean.

**Abbreviations:** OLCs: Oligodendrocyte-like cells; NBs: neuroblasts; TAPs: transient amplifying progenitors; mTAPs: mitotic TAPs; qNSCs: quiescent neural stem cells; aNSCs: activated NSCs.

**Figure S7.**
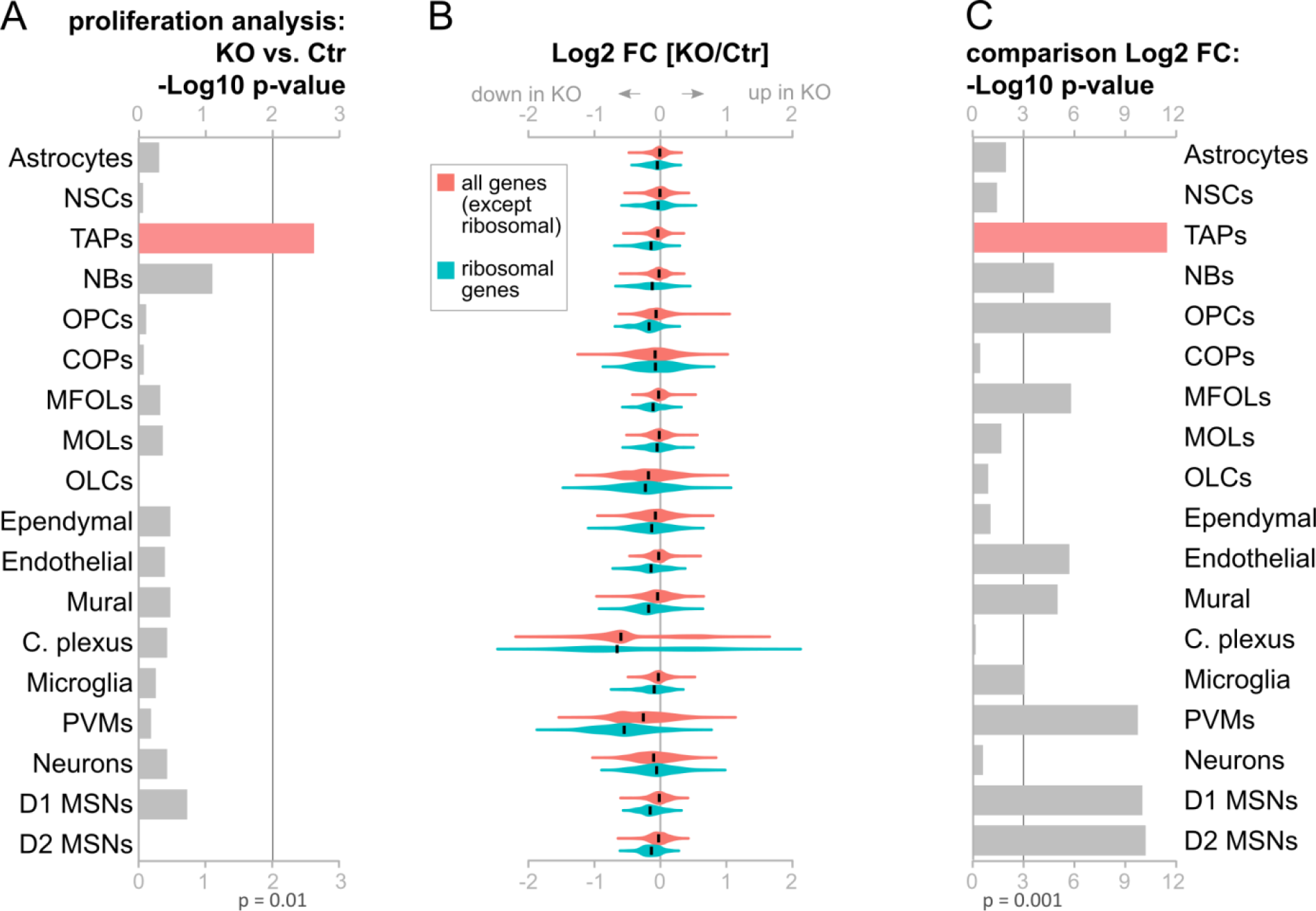
Proliferation is significantly reduced in TAPs, and ribosomal genes are decreased in LRP2-deficient SVZ. Relate to Figure 4 and 5. (A) P-values of the proliferation analysis for all cell types. Statistical significance of data was determined using Kolmogorov-Smirnov test on the cumulative fraction distribution of KO vs. Ctr cells scored for cell cycle. The p-value for TAPs is highlighted in red. (B) Distribution of KO vs. Ctr expression of ribosomal (red) and all remaining genes (teal). Expression cut off ≥ 0.5 log2 (mean normalized UMI count) Ctr cells. (C) P-values for the comparison of the distribution of ribosomal and all remaining genes in KO vs. Ctr cells. In 10 out of 18 cell clusters, ribosomal genes are significantly lower expressed as all remaining genes in KO compared to Ctr cells (p-value < 0.001). TAPs (highlighted in red) had the smallest p-value. Statistical significance of data was determined using unpaired student’s t test.

**Abbreviations:** FC: Fold Change; all other abbreviations as in Figure S5.

**Table S1: List of significantly upregulated genes in all identified cell types. Related to Figure 2.**

**Table S2: List of significantly up- and downregulated genes in NSCs compared to niche astrocytes. Related to Figure 2.**

**Table S3: List of significantly upregulated genes in subclusters. Related to Figure 3.**

**Table S4: Differentially expressed genes per cell cluster of LRP2-deficient brains. Related to Figure 5.**

